# A spatio-temporally constrained gene regulatory network directed by PBX1/2 acquires limb patterning specificity via HAND2

**DOI:** 10.1101/2022.03.08.483529

**Authors:** Marta Losa, Iros Barozzi, Marco Osterwalder, Peyman Zarrineh, Jean Denis Benazet, Brandon Chacon, Ausra Girdziusaite, Angela Morabito, Jianjian Zhu, Susan Mackem, Terence D. Capellini, Nicoletta Bobola, Diane Dickel, Aimee Zuniga, Axel Visel, Rolf Zeller, Licia Selleri

**Affiliations:** Program in Craniofacial Biology, Institute for Human Genetics, Eli and Edythe Broad Center of Regeneration Medicine and Stem Cell Research, Department of Orofacial Sciences and Department of Anatomy, University of California at San Francisco, San Francisco, California, USA; Center for Cancer Research, Medical University of Vienna, Vienna, Austria; Environmental Genomics and Systems Biology Division, Lawrence Berkeley National Laboratory, Berkeley, CA, USA; Department for Biomedical Research (DBMR), University of Bern, Bern, Switzerland; Department of Cardiology, Bern University Hospital, Bern, Switzerland; School of Medical Sciences, University of Manchester, Manchester, United Kingdom; Developmental Genetics, Department Biomedicine, University of Basel, Basel, Switzerland; Cancer and Developmental Biology Laboratory, Center for Cancer Research, National Cancer Institute, Frederick, MD, USA; Department of Human Evolutionary Biology, Harvard University, Cambridge, MA; Broad Institute of Harvard and MIT, Cambridge, MA, USA; US Department of Energy Joint Genome Institute, Lawrence Berkeley National Laboratory, Berkeley, CA 94720,USA; School of Natural Sciences, University of California, Merced, Merced, CA 95343, USA

**Author notes:** Equal contributions (Marta Losa, Iros Barozzi). Co-senior authors (Rolf Zeller, Licia Selleri).

## Abstract

During development cell fates are specified by tightly controlled gene expression programs. PBX TALE transcription factors control gene regulatory networks (GRN) that direct vertebrate tissue patterning and organ morphogenesis. How PBX1/2 proteins acquire context-specific functions, despite widespread embryonic expression of *Pbx1/2*, remains elusive. In mouse limb buds, mesenchymal-specific loss of PBX1/2 or of the transcriptional regulator HAND2 results in similar phenotypes, suggesting that PBX1/2- and HAND2-dependent programs converge to control limb development. To investigate this scenario, we combined tissue-specific and temporally-controlled mutagenesis with multi-omics approaches using the murine hindlimb bud as a model. We reconstructed a GRN collaboratively directed by PBX1/2 and HAND2, demonstrating that *Pbx1*-*Hand2* genetically interact *in vivo* during hindlimb patterning, with PBX1 concomitantly acting as an upstream regulator of *Hand2*. At organismal-level resolution the GRN is active within restricted subsets of posterior-proximal hindlimb mesenchymal cells, wherein *Pbx1/2* and *Hand2* are co-expressed with their target genes. Genome-wide profiling of PBX1 binding across multiple tissues further revealed that HAND2 selects a subset of PBX-bound regions to impart limb patterning functionality. This research elucidates mechanisms underlying limb bud-specific functions by PBX1/2, while informing general principles by which promiscuous transcription factors cooperate with select cofactors to instruct distinct developmental programs.

## INTRODUCTION

Through the years, the limb has provided an excellent model for understanding the genetic principles that control development, evolution, and congenital disease (Royle et al. 2021; Zuniga and Zeller 2020). Numerous genetic studies in the mouse have led to significant insights into the genes and molecular pathways that direct limb bud patterning and morphogenesis. Our research has established that the gene family encoding PBX1/2/3 homeodomain transcription factors of the TALE superclass are essential regulators of limb development (Moens and Selleri 2006; Capellini et al. 2011b). PBX homeoproteins execute hierarchical, overlapping, and iterative functions during limb development; they are essential in determining limb bud positioning and formation, limb axes establishment, as well as patterning and morphogenesis of most skeletal elements of the limb and girdle (Capellini et al. 2011b, 2008, 2006, 2011a). In particular, PBX homeoproteins control effectors of limb and girdle development, including genes encoding the SHH morphogen in the limb posterior mesenchyme (Capellini et al. 2006) and transcriptional regulators such as ALX1 in pre-scapular domains (Capellini et al. 2010).

We demonstrated that PBX homeoproteins do not act solely as cofactors for HOX proteins in limb buds (Moens and Selleri 2006), but also execute upstream control of 5' *HoxA/D* gene expression (Capellini et al. 2006). In particular, homozygous loss of *Pbx1* (*Pbx1^−/−^*) causes malformations of girdles and proximal stylopod skeletal structures (Selleri et al. 2001), while homozygous inactivation of *Pbx2* or *Pbx3* does not yield limb skeletal phenotypes (Selleri et al. 2004; Rhee et al. 2004). In contrast, compound constitutive loss-of-function of *Pbx1/2* results in multiple developmental abnormalities, including striking distal limb defects with loss of posterior digits (Capellini et al. 2006), and exacerbates the proximal limb phenotypes reported in *Pbx1^−/−^* embryos (Selleri et al. 2001). Given that the mouse hindlimb bud lacks detectable *Pbx3* expression (di Giacomo et al. 2006) and that *Pbx4*, the last known member of the *Pbx* family, is not expressed during limb bud development (Wagner et al. 2001), compound loss of *Pbx1/2* in the developing hindlimb achieves a PBX-null state. Accordingly, *Pbx1/2* mutant hindlimbs exhibit more pronounced phenotypes than those observed in forelimbs, which express high levels of *Pbx3* (di Giacomo et al. 2006). However, compound constitutive homozygous deletion of *Pbx1* and *Pbx2* results in embryonic lethality by gestational day (E)10.5 (Capellini et al. 2006) due to striking cardiovascular defects (Stankunas et al. 2008), which prevents a mechanistic understanding of their key functions as transcriptional regulators of limb bud development.

The establishment of *cis*-regulatory landscapes and gene regulatory networks (GRNs) that control limb patterning and outgrowth has provided an additional layer to our understanding of this archetypical developmental process (Bolt and Duboule 2020). For example, dissection of the chromatin organization and regulation of the murine *Hox* gene clusters (Amândio et al. 2021; Fabre et al. 2018) has contributed mechanistic insight into the critical roles of *Hox* genes in setting up regional identities along the body axis of bilaterians and during limb bud patterning (Darbellay and Duboule 2016). Specifically, expression of all four *Hox9* paralogs is required for anterior-posterior (AP) polarization of the limb bud and initiation of SHH signaling (Xu and Wellik 2011). Of note, our studies have shown that establishment of *Shh* expression in the limb zone of polarizing activity (ZPA) requires PBX1/2 (Capellini et al. 2006) and the bHLH transcription factor HAND2 (Galli et al. 2010). In particular, *Hand2* is part of a GRN that antagonizes *Gli3* expression to establish AP asymmetry and pentadactyly (Galli et al. 2010; te Welscher et al. 2002). Establishment of the posterior domain of *Hand2* expression is not only dependent on its mutual antagonistic interaction with *Gli3* and early-wave *Hox* genes, but also on HOX-interacting MEIS1/2 transcription factors that bind to a *Hand2* limb enhancer (Delgado et al. 2021). Identification of HAND2 targets has uncovered a GRN of regulators that control compartmentalization of the limb bud mesenchyme, emphasizing an evolutionary conserved role of HAND2 in orchestrating early limb development upstream of SHH (Osterwalder et al. 2014). Lastly, the genetic dissection of the *Gremlin1* (*Grem1*) *cis*-regulatory landscape, which directs the spatio-temporal expression of *Grem1* in limb buds, has provided molecular insight into the robustness and evolutionary plasticity of distal limb and digit development (Malkmus et al. 2021).

While it was established that both PBX1/2 and HAND2 independently activate posterior *Shh* expression by directly interacting with its distal limb enhancer (called ZRS) (Sagai et al. 2005; Capellini et al. 2006; Galli et al. 2010), it is unknown whether their functions converge on regulating SHH signaling. If this is the case, potential genetic interactions between PBX1/2 and HAND2 and GRNs jointly controlled by these factors remain elusive. Here, we combined tissue-specific and temporally controlled gene inactivation in mouse embryonic hindlimb buds with a variety of transgenic and multi-omics approaches using dissected limb buds to reconstruct a multi-layered GRN of limb regulators that is collaboratively directed by PBX1/2 and HAND2. We further established that the genetic interaction of *Pbx1* with *Hand2* in the limb is required for normal hindlimb bud development and that PBX1 acts also as an upstream regulator of *Hand2* expression. Lastly, by comparing PBX genome-wide occupancy *in vivo* across multiple embryonic tissues, we uncovered that PBX transcription factors indiscriminately occupy a vast pool of common genomic loci during development. However, promiscuous PBX binding gains restrained functionality via cooperative interactions with select cofactors that promote tissue specificity. Together, these studies enabled the reconstruction of a spatio-temporally constrained GRN at superior organismal- and tissue-level resolution. The PBX1/2-directed GRN acquires early limb patterning functions via HAND2, a critical regulator of limb AP asymmetry and pentadactyly.

## RESULTS

### Requirement of PBX1 and PBX2 in limb bud mesenchyme during the onset of limb bud patterning and outgrowth

Given the dominant role of PBX1 among PBX family members during hindlimb development (Capellini et al. 2006), we examined its spatio-temporal distribution from formation of the hindlimb bud (E9.0) to digit development (E12.0). Following its expression in the lateral plate mesoderm (E8.0, Capellini et al. 2006), PBX1 was detected in most limb mesenchymal progenitors and in the apical ectodermal ridge (AER) during the onset of hindlimb bud development (E9.0-10.5, Fig.1A-C’), while it was restricted to the proximal-most mesenchyme by E11.0 until E12.0 (Fig. S1A-C). To circumvent early embryonic lethality of *Pbx1/2* compound constitutive null embryos and to decipher tissue-specific *Pbx* functions during limb development, we conditionally inactivated *Pbx1* (Koss et al. 2012) on a *Pbx2*-deficient background (Selleri et al. 2004). This genetic approach enabled us to obtain a *Pbx*-null state in the hindlimb bud, where *Pbx3* is not expressed at detectable levels (di Giacomo et al. 2006). AER-specific deletion of *Pbx1* using a *Msx2Cre* deleter line (Liu et al. 1994; Sun et al. 2000) did not cause any limb skeletal abnormalities (Fig. S1D-G, Supplementary Table 1) and was verified by immunohistochemistry (Fig. S1H-I’). These genetic experiments demonstrate that PBX1 is dispensable for AER formation and maintenance.

**Figure 1.**
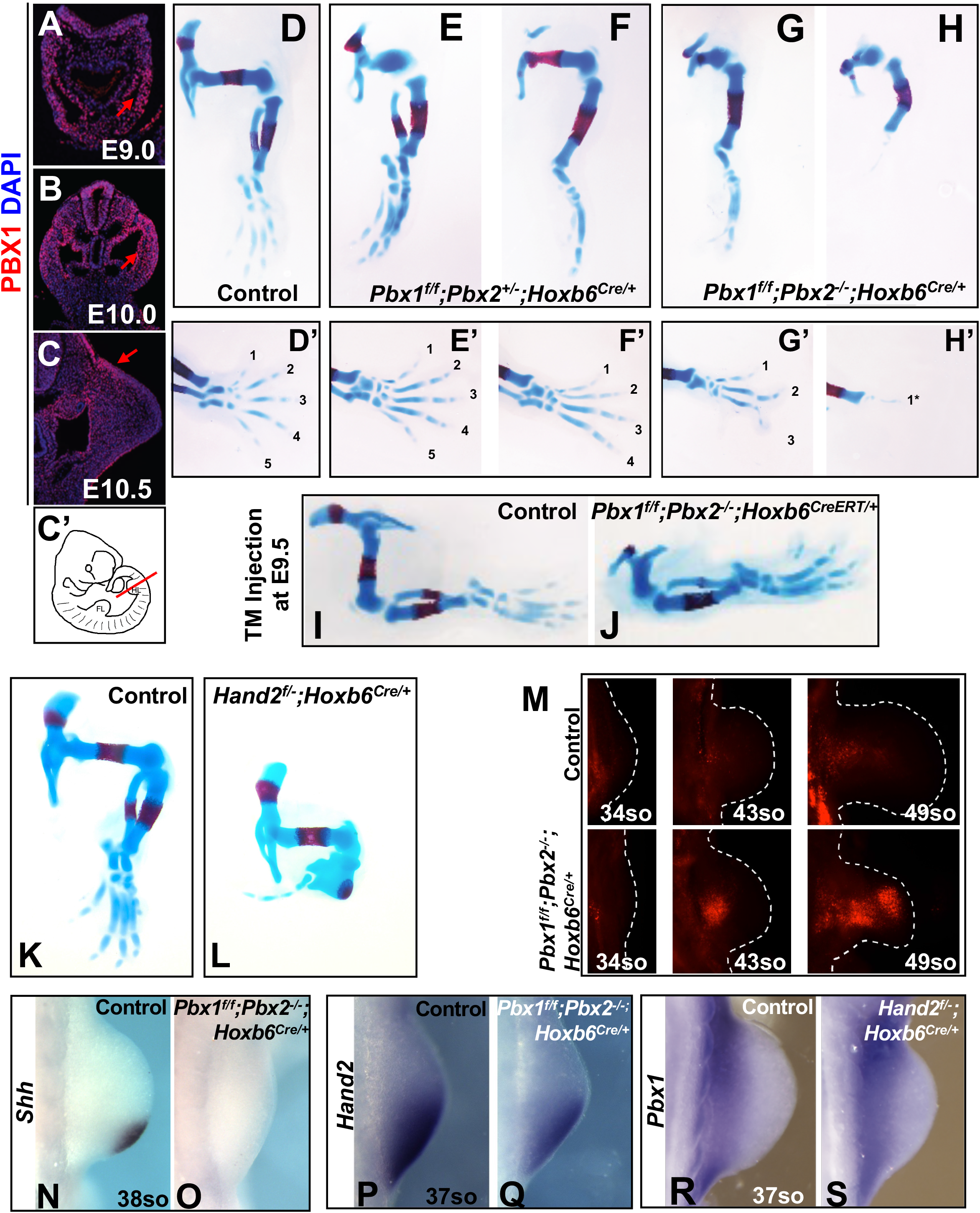
PBX1/2 are required in the mesenchyme for hindlimb patterning before E9.5. (A-C’) IF showing PBX1 protein (red) localization in developing murine hindlimbs from E9.0 to E10.5. DAPI labels nuclei (blue); C’ shows plane of section through hindlimb bud in E10.5 mouse embryo. (D-H) E14.5 skeletal preparations of mutant hindlimbs with tissue-specific deletion of *Pbx1* on a *Pbx2*-deficient background using a *Hoxb6Cre* deleter line (referred in text as *Pbx1cKO^Mes^*;*Pbx2^+/−^* and *Pbx1cKO^Mes^*;*Pbx2^−/−^*); cartilage and bone visualized by Alcian Blue and Alizarin Red staining, respectively. Digits shown from anterior (digit 1) to posterior (digit 5). (I,J) Representative images of *Pbx1^f/f^;Pbx2^−/−^;Hoxb6^CreERT/+^* hindlimb skeletal phenotype compared to *Pbx1^f/f^;Pbx2^−/−^* Control. Ratio of tibia/femur length in mutant embryos *versus* controls quantified in Suppl. Figure 1 (Panel J). (K,L) E14.5 skeletal preparations of mutant hindlimbs with tissue-specific deletion of *Hand2* using the *Hoxb6Cre* deleter (*Hand2^f/−^;Hoxb6^Cre/+^*). (M) Detection of apoptotic cells by LysoTracker Red. *Pbx1^f/f^;Pbx2^−/−^;HoxB6^Cre/+^* hindlimb buds and respective controls at E10.25 (34 somites), E11.0 (43 somites), and E11.5 (48 somites). (N-S) Whole mount *in situ* hybridization (WISH) of *Shh* (N,O) and *Hand2* (P,Q) in E10.75 *Pbx1cKO^Mes^*;*Pbx2^−/−^* hindlimbs *versus* controls. (R,S) WISH of *Pbx1* in E10.75 *Hand2cKO^Mes^* hindlimb *versus* controls. In all panels, hindlimb buds oriented with anterior to the top and posterior to the bottom.

To investigate PBX functions during early limb bud development, the *Hoxb6Cre* deleter line expressing the *Cre* recombinase under the control of the *Hoxb6* promoter (Lowe et al. 2000) was used for conditional gene inactivation in the lateral plate mesoderm, posterior forelimb and entire hindlimb bud mesenchyme from limb initiation onward. Mesenchyme-specific deletion of *Pbx1* on a *Pbx2*-deficient background resulted in drastic limb defects (Fig. 1D-H) that phenocopy the limb skeletal abnormalities of constitutive compound *Pbx1/2* mutants (Capellini et al. 2006). Although the phenotypes of *Pbx1^f/f^;Pbx2^+/−^;Hoxb6^Cre/+^* (hereafter *Pbx1cKO^Mes^*;*Pbx2*^+/−^) (Fig. 1E,F) and *Pbx1^f/f^;Pbx2^−/−^;Hoxb6^Cre/+^* (hereafter *Pbx1*cKO^Mes^;*Pbx2*^−/−^) hindlimbs (Fig. 1G,H) displayed variable penetrance, the pelvic girdle was dysmorphic and the proximal as well as distal elements were malformed, hypoplastic, or absent in all mutant embryos at E14.5 (Supplementary Table 1). As expected, phenotypes were more severe when both *Pbx1* and *Pbx2* were inactivated, varying from hypoplastic autopodia (Fig. 1G’) to complete autopod agenesis (Fig. 1H’). To assess the temporal requirement of *Pbx1* during hindlimb bud development, we used an inducible *Hoxb6Cre* (*Hoxb6^CreERT^*) line (Nguyen et al. 2009). Tamoxifen injections at E8.5, E9.5, E10.0 and E10.5, respectively, showed that *Pbx1/2* are required in the mesenchyme for hindlimb patterning prior to E9.5 (Fig. 1I,J and Fig. S1J), as their inactivation at or before this time-point resulted in hindlimb abnormalities that mimicked those observed using the *Hoxb6Cre* allele. In contrast, hindlimbs developed normally when tamoxifen was administered at or after E10.0. Altogether, concomitant inactivation of *Pbx1/2* with different *Cre* deleter lines (Fig. 1D-J, S1D-G and S1J) demonstrates that PBX1/2 are dispensable in the AER and are required in the hindlimb mesenchyme at least until E9.5, i.e. during hindlimb bud initiation and early patterning.

### Early mesenchymal-specific deletion of *Pbx1/Pbx2* or *Hand2* results in similar skeletal, cellular, and molecular alterations

The hindlimb skeletal phenotypes of E14.5 *Pbx1cKO^Mes^*;*Pbx2*^−/−^ mouse embryos are similar to those reported in embryos with mesenchymal loss of *Hand2* (hereafter *Hand2cKO^Mes^*) (Galli et al. 2010; Osterwalder et al. 2014) (Fig. 1K,L) and constitutive loss of *Shh (Shh^−/−^*; Chiang et al. 2001). Given the striking similarities of the distal phenotypes in these mutant limbs, we sought to investigate the cellular and molecular alterations underlying the respective skeletal defects. Limb bud mesenchymal apoptosis (Fernández-Terán et al. 2006) was not increased in *Pbx1cKO^Mes^*;*Pbx2*^−/−^ hindlimb buds in comparison to controls at early stages (34 somites, Fig. 1M). However, later in development, apoptosis of mesenchymal cells was significantly increased in the distal-anterior hindlimb bud mesenchyme of *Pbx1cKO^Mes^;Pbx2^−/−^* embryos (43 and 49 somites, Fig. 1M). These findings were consistent with increased mesenchymal apoptosis in *Shh-* and *Hand2*-loss-of-function limb buds (Chiang et al. 2001; Galli et al. 2010; Charite et al. 2000). Interestingly, *Pbx1/2*-deficient hindlimb buds expressed negligible levels of *Shh* (Capellini et al. 2006). RNA *in situ* hybridization corroborated complete loss of *Shh* also in *Pbx1cKO^Mes^;Pbx2^−/−^* hindlimbs (Fig. 1N,O), as was the case in *Hand2cKO^Mes^* hindlimb buds (Galli et al. 2010; Osterwalder et al. 2014). These results demonstrate that the striking similarities observed in the distal limb skeletal phenotypes of *Pbx1cKO^Mes^*;*Pbx2^−/−^* and *Hand2cKO^Mes^* embryos are underpinned by similar cellular and molecular alterations.

Given these similarities, we asked whether *Pbx1/2* and *Hand2* converge in orchestrating a shared GRN essential for limb bud patterning and morphogenesis beyond their independent requirements in *Shh* regulation. First, to assess a potential epistatic hierarchy, we examined the spatial expression of *Hand2* transcripts in *Pbx1cKO^Mes^;Pbx2^−/−^* hindlimb buds. Compared to controls, *Hand2* transcript levels were reduced in the proximal-most mesenchyme of *Pbx1cKO^Mes^;Pbx2^−/−^* developing hindlimbs (Fig. 1P,Q), as reported for *Pbx1/2* constitutive mutants (Capellini et al. 2006). In contrast, *Pbx1* spatial expression was not altered in *Hand2cKO^Mes^* hindlimb buds (Fig. 1R,S). Altogether, these findings corroborate previous reports that both PBX1/2 and HAND2 are essential to establish *Shh* expression in the posterior limb bud mesenchyme. They further suggest that *Pbx1* contributes to the regulation of *Hand2* expression, likely within a GRN that orchestrates the onset of limb bud patterning and outgrowth.

### *Pbx1/2* and *Hand2* are co-expressed in a subset of posterior-proximal hindlimb bud mesenchymal cells

To investigate the genetic hierarchy, individual contributions, and convergence of PBX and HAND2 in regulating downstream limb GRNs, we combined mouse genetics with genome-wide transcriptome and epigenome profiling of dissected hindlimb buds (Fig. 2A). We isolated hindlimb buds from wild-type mouse embryos at E10.5 (36-38 somites) and performed single cell (sc) RNAseq analysis (Cao et al. 2019) to disentangle the cellular heterogeneity of early hindlimb development and assess potential co-expression of *Pbx1/2* and *Hand2* transcripts at single-cell resolution (Fig. 2B-F and Fig. S2,S3). Unsupervised clustering and limb bud marker gene analysis led to the identification of six main cell populations within E10.5 hindlimbs, including mesenchymal cells (*Prrx1+* and *Meis2+*); epithelial cells and AER (*Epcam+* and *Wnt6+*); erythrocytes (*Hba-a2+* and *Klf1+*); endothelial cells (*Pecam1+* and *Emcn+*); phagocytic cells (*Fcer1g+* and *Spi1+*); and pluripotent progenitor cells (*Sox2+* and *Pou5f1+*) (Fig. 2B,C, Fig. S2A,B, and S3A-F; Kelly et al. 2020; Desanlis et al. 2020). *Pbx1* and *Pbx2* were detected in both the mesenchymal and epithelial cell populations, although their expression was predominant in mesenchymal cells, with *Pbx1* being overall more abundant than *Pbx2* (Fig. 2D and Fig. S2C). In contrast, *Hand2* expression was restricted to a subset of posterior mesenchymal cells (Fig. 2E, Galli et al. 2010; Osterwalder et al. 2014). Additional analyses confirmed that *Hand2* is co-expressed with *Pbx1* and *Pbx2* in a fraction of mesenchymal cells (orange and red, Fig. 2F). These mesenchymal subpopulations were characterized by high levels of *Tbx3* and *Isl1* and low levels of *Lhx9* and *Lhx2* transcripts, which identify them as posterior-proximal hindlimb bud mesenchyme (Fig. S2B and S3G). In agreement, these subpopulations lacked anterior-proximal markers such as *Irx3* and *Irx5* (Fig. S3G). The expression of *HoxA* and *HoxD* genes further confirmed the spatial assignment of these cells to the posterior-proximal mesenchyme (Fig. S3G). In addition, immunofluorescence (IF) analysis of hindlimb bud sections revealed co-localization of PBX1 and HAND2 proteins in the posterior mesenchyme (Fig. 2G-J’’). PBX1 was present in most mesenchymal cell nuclei, although at lower levels in the distal hindlimb mesenchyme, while HAND2 was restricted to the posterior-proximal mesenchyme. These results show that nuclear PBX1 and HAND2 transcription factors co-localize in the same posterior-proximal mesenchymal cells in hindlimb buds.

**Figure 2.**
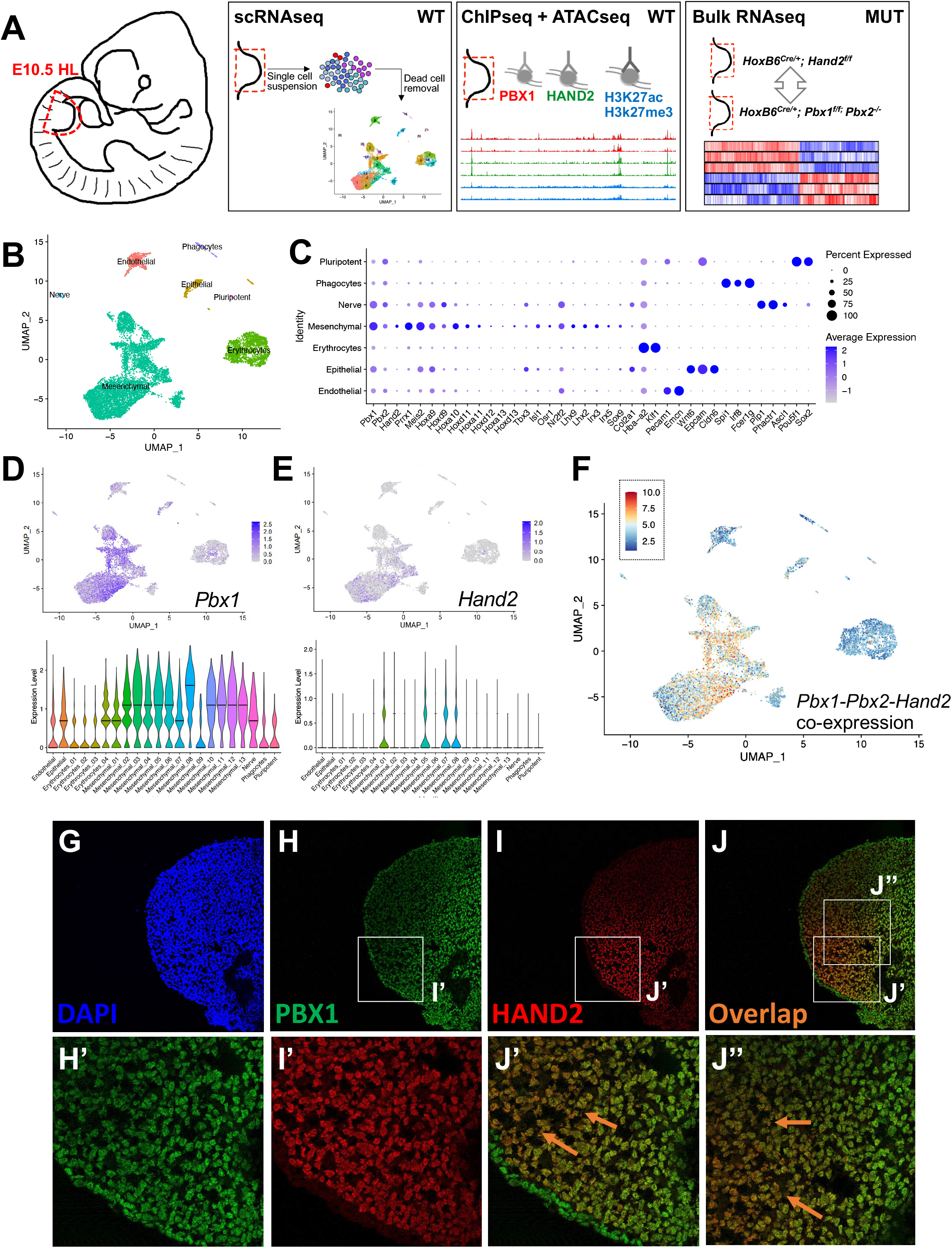
Identification of hindlimb bud mesenchymal subpopulations co-expressing *Pbx1/2-Hand2*. (A) Workflow of the genome-wide approaches performed in parallel in this study. (B) Uniform manifold approximation and projection (UMAP) representation of high-quality 9,859 hindlimb cells at E10.5-10.75, defined by scRNAseq. (C) Dot plot of top differentially expressed markers for each embryonic hindlimb population. Size of each dot represents proportion of cells within a given population that expresses the gene; intensity of color indicates average level of gene expression in indicated population. (D,E) UMAPs (upper) and violin plots (lower) showing normalized expression patterns for *Pbx1* and *Hand2*. Y axes in violin plots are at saturation and differ in scale. (F) UMAP highlighting co-expression of *Pbx1-Pbx2*-*Hand2.* (G-J’) IF of PBX1 (green) and HAND2 (red) proteins in posterior-proximal hindlimb mesenchyme. Higher magnification shows co-localization (orange) in H’, I’ J’ and J’’ insets (orange arrows; double positive cells). DAPI (blue) labels nuclei.

### PBX and HAND2 regulate a GRN that controls early hindlimb bud development

To identify unique and shared transcriptional targets of PBX and HAND2 and their epigenetic landscapes in mouse hindlimb buds, we used chromatin immunoprecipitation followed by sequencing (ChIPseq, Amin et al. 2015; Losa et al. 2017), in combination with ATACseq (genome-wide assay for transposase-accessible chromatin using sequencing, Buenrostro et al. 2013) in E10.5 hindlimb buds (middle panel, Fig. 2A). From all datasets obtained, only the peaks detected reproducibly across two replicates were selected for further analysis (Supplementary Table 2; see Methods). Replicated peaks for PBX1 and HAND2 were merged into one set of 32,691 putative regulatory elements: 6,157 bound by both transcription factors; 4,536 bound only by HAND2; and 21,998 bound only by PBX1 (Figure 3A,B and S4A-C). The majority of HAND2 peaks were located distal to the transcription start site (TSS, Osterwalder et al. 2014) of annotated genes, while PBX1 peaks were distributed between promoters and TSS-distal regions (Fig. 3A,B). Regions co-bound by both PBX1 and HAND2 predominantly associated with distal elements, including both intergenic and intragenic regions (Fig. 3A). Remarkably, sites co-bound by PBX1 and HAND2 were located in regions of higher chromatin accessibility (Fig. 3C; *p*-value < 2.2e-16, Kruskal-Wallis Test) and had greater enrichment values for transcription factor-binding (Fig. 3D,E; *p*-value < 2.2e-16, Mann-Whitney Tests) than sites bound by only one of the two factors. Genomic Regions Enrichment Annotation (GREAT, McLean et al. 2010) analysis revealed that the peaks bound by both PBX1 and HAND2 were mostly associated with genes known to regulate limb bud and/or skeletal development (Fig. 3F,G), while this was not the case for regions bound only by one transcription factor (Fig. S4D-G).

**Figure 3.**
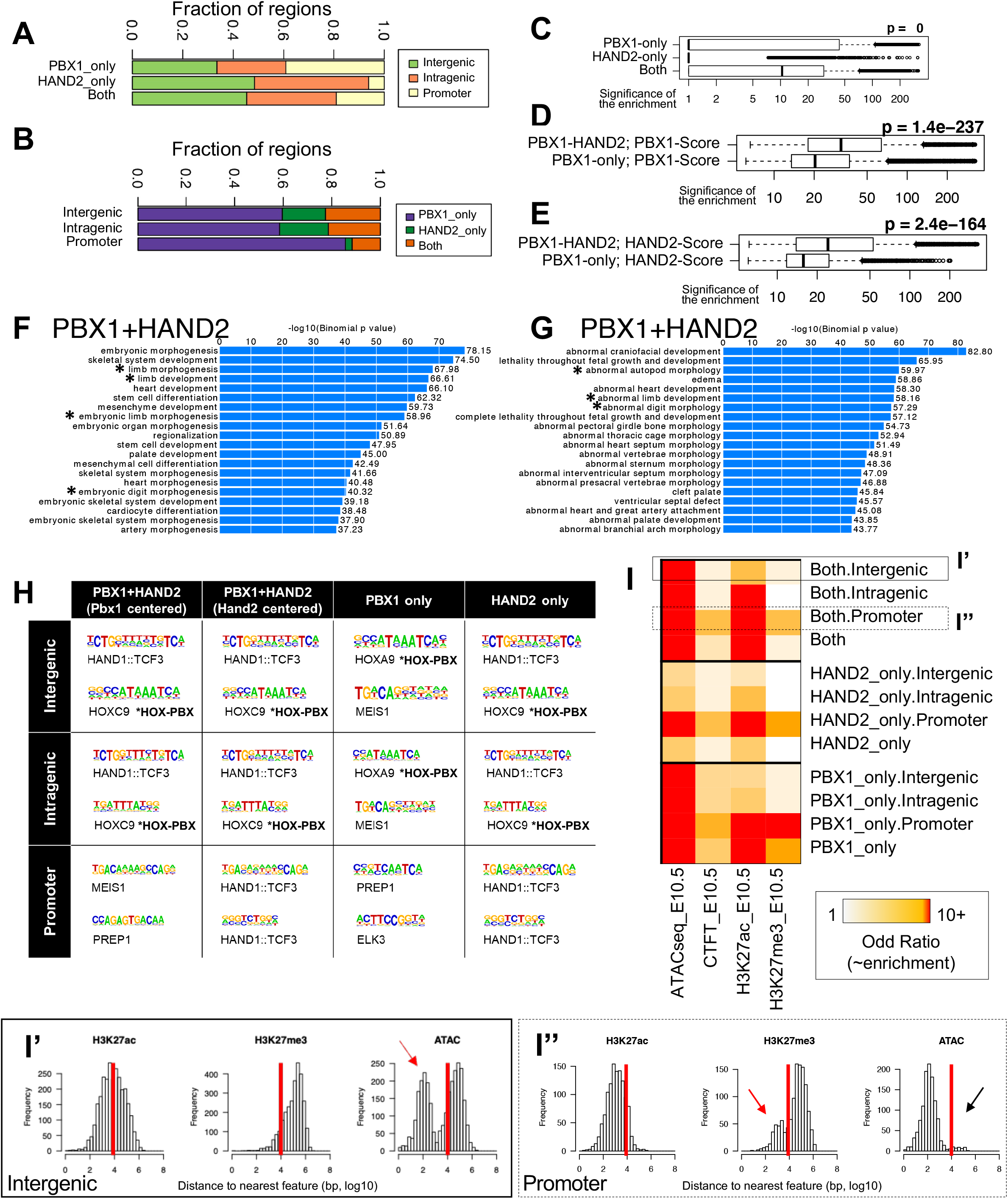
PBX1 and HAND2 control a shared gene regulatory network that drives early hindlimb development. (A) Distribution of the proportion of PBX1-only, HAND2-only or PBX1-HAND2 co-bound peaks relative to gene intergenic, promoter, or intragenic regions. (B) Similar to (A); showing proportion of intergenic, promoter, or intragenic regions relative to PBX1-only, HAND2-only, or PBX1-HAND2 co-bound peaks. (C) Assessment of chromatin accessibility for PBX1-only and HAND2-only bound peaks *versus* PBX1-HAND2 co-bound peaks. X axis shows −log10(p-value) of ATACseq enrichment, on logarithmic scale. (D-E) Assessment of enrichment values for PBX1-only and HAND2-only bound peaks *versus* PBX1-HAND2 co-bound peaks. X axis shows the −log10(*p*-value) of ChIPseq enrichment, on logarithmic scale. (F,G) Top over-represented biological processes (F) and mouse phenotypes (G) associated with PBX1-HAND2 co-bound peaks. X axes show −log10 of binomial (uncorrected) *p*-values. Limb-related categories highlighted by asterisk*. (H) Top two most significantly enriched motifs identified by *de novo* motif enrichment analysis of peaks co-bound by PBX1-HAND2, PBX1-only, or HAND2-only (*q*-value <= 0.05). Identity of the most similar annotated motif indicated below each logo. (I) Heatmap summarizing enrichment (as odds ratio of observed *versus* expected fraction of peaks) of indicated chromatin features (columns) at peaks co-bound by PBX1-HAND2 (both), PBX1-only, or HAND2-only (rows). (I’,I’’) Distribution of H3K27ac/H3K27me3 or ATACseq peaks relative to distance to nearest peak bound by both PBX1-HAND2 in intergenic sites (I’) or promoters (I’’).

*De novo* motif discovery (using HOMER, Heinz et al. 2010) determined whether both PBX1 and HAND2, or only one of them, interacted directly with DNA or as part of transcriptional complexes (Fig. 3H). The HAND2 (annotated by similarity to HAND1::TCF3) and HOX-PBX (TGATTNTT; annotated to HOXC9) motifs were identified as the top two most-enriched motifs in shared peaks, whether they were centered on the PBX1 peak or the HAND2 peak (Fig. 3H; p-value < 0.05). Interestingly, the HOX-PBX motif was still the second most enriched sequence motif in peaks bound by HAND2 even in the absence of PBX1 binding. In contrast, PBX1 bound on its own with other TALE factors (MEIS/PREP) at promoter regions and distal elements (Fig. 3H; *p*-value < 0.05). Of note, the majority of genes associated with PBX1-specific peaks function in developmental processes other than limb morphogenesis (Fig. S4F). Moreover, motif analysis using a large set of published binding preferences (Heinz et al. 2010) revealed significant associations of the PBX1-only peaks with binding of other homeobox transcription factors (HOX, MEIS, PDX2, LHX2) at TSS-distal sites (Fig. S4H; *p*-value < 1e-5; HOMER). These results point to the convergence of PBX1 and HAND2 on *cis*-regulatory modules within a GRN in the posterior-proximal limb mesenchyme that orchestrates limb and digit patterning (asterisks, Fig. 3F,G).

Next, we integrated our sets of PBX1 and HAND2 replicated peaks with the H3K27ac (associated with active enhancer marks) and H3K27me3 (associated with repressive enhancer marks) ChIPseqs (Lee and Young 2013) and the ATACseq profiles (denoting open chromatin) that we generated from E10.5 hindlimb buds. In addition, we intersected these data sets with published CTCF binding profiles (Fig. 3I, DeMare et al. 2013). Our analyses revealed that: 1) Intergenic, intragenic and promoter regions co-bound by PBX1 and HAND2 associate with accessible chromatin and H3K27ac enrichment (Fig. 3I’,I’’); 2) regions bound by PBX1 only are significantly more accessible and more enriched with H3K27ac than regions bound by HAND2 only (Fig. 3I and Fig. S5A-D); 3) no significant associations are present with genomic regions marked by H3K27me3 repressive marks, with the exception of a fraction of promoter regions bound by PBX1 (red arrow, Fig. S5D), which points to potential bivalent promoters; and 4) as expected, CTCF binding is not associated with any of the groups analyzed. Together, these analyses indicate that PBX1 and HAND2 interact with a common set of candidate *cis*-regulatory modules (CRMs) in accessible and active chromatin within genomic landscapes of genes that regulate limb development.

### PBX1 directly regulates *Hand2* expression via specific CRMs active in the hindlimb bud mesenchyme

In light of the genetic evidence indicating that *Pbx1/2* regulate *Hand2* expression in the proximal-posterior limb bud mesenchyme (Fig. 1P,Q and Fig. 2D-F), we set out to identify the complement of limb enhancers critical for this process. To this end, we screened the *Hand2* TAD (Bonev et al. 2017) by intersecting our PBX1, HAND2, H3K27ac ChIPseq and ATACseq datasets (Fig. 4A) (see Methods). This analysis uncovered 17 PBX1-bound putative CRMs with predicted limb enhancer activity, including 3 previously identified candidate limb enhancers (mm1687, mm1688, and mm1689) as validated by transgenic *LacZ* reporter assays at E11.5 (Fig. 4B,C,E, Supplementary Table 3; Monti et al. 2017). A similar strategy was used to analyze the remaining putative PBX1-bound limb enhancer elements (n=14) in mouse embryos at E10.5 or E11.5. These assays revealed the presence of 7 additional tissue-specific enhancers in the *Hand2* TAD with activity in limb buds and/or other embryonic compartments (Fig. 4D,F,G and Fig. S6; Supplementary Table 3). Of these, 3 showed reproducible limb activities in *Hand2* expressing domains, expanding the number of known *bona fide* embryonic limb enhancers in the *Hand2* locus to 6 in total, all of which were active at E10.5 and E11.5 (Fig. 4B-G and Fig. S6). However, only 2 of these enhancers, located 337kb (mm1689) and 226kb (mm1828) upstream of the *Hand2* TSS, respectively, displayed activity restricted to the posterior hindlimb bud mesenchyme at E10.5 (Fig. 4E,F; dashed rectangles) overlapping the endogenous *Hand2* expression domain. Interestingly, it was recently reported that deletion of mm1689 in the mouse results in loss of *Hand2* expression in limb buds (Delgado et al. 2021). To demonstrate that the activity of this CRM is regulated by PBX, we mutagenized all PBX and PBX-HOX binding sites within this element (Fig. 4H and Supplementary Methods), disrupting the conserved core of the binding motifs (Moens and Selleri 2006; Selleri et al. 2019). We then assayed the mutagenized (MUT) and wild-type (WT) versions of this enhancer using a CRISPR/Cas9-mediated site-specific transgenic mouse assay termed enSERT (Kvon et al. 2020). Activity of the mm1689 enhancer was completely lost with high reproducibility in its MUT version (n=9) compared to WT (n=2) (Fig 4H). Together, our results indicate that in early hindlimb buds the activity of CRMs in the *Hand2* genomic landscape critically depends on intact PBX binding sites.

**Figure 4.**
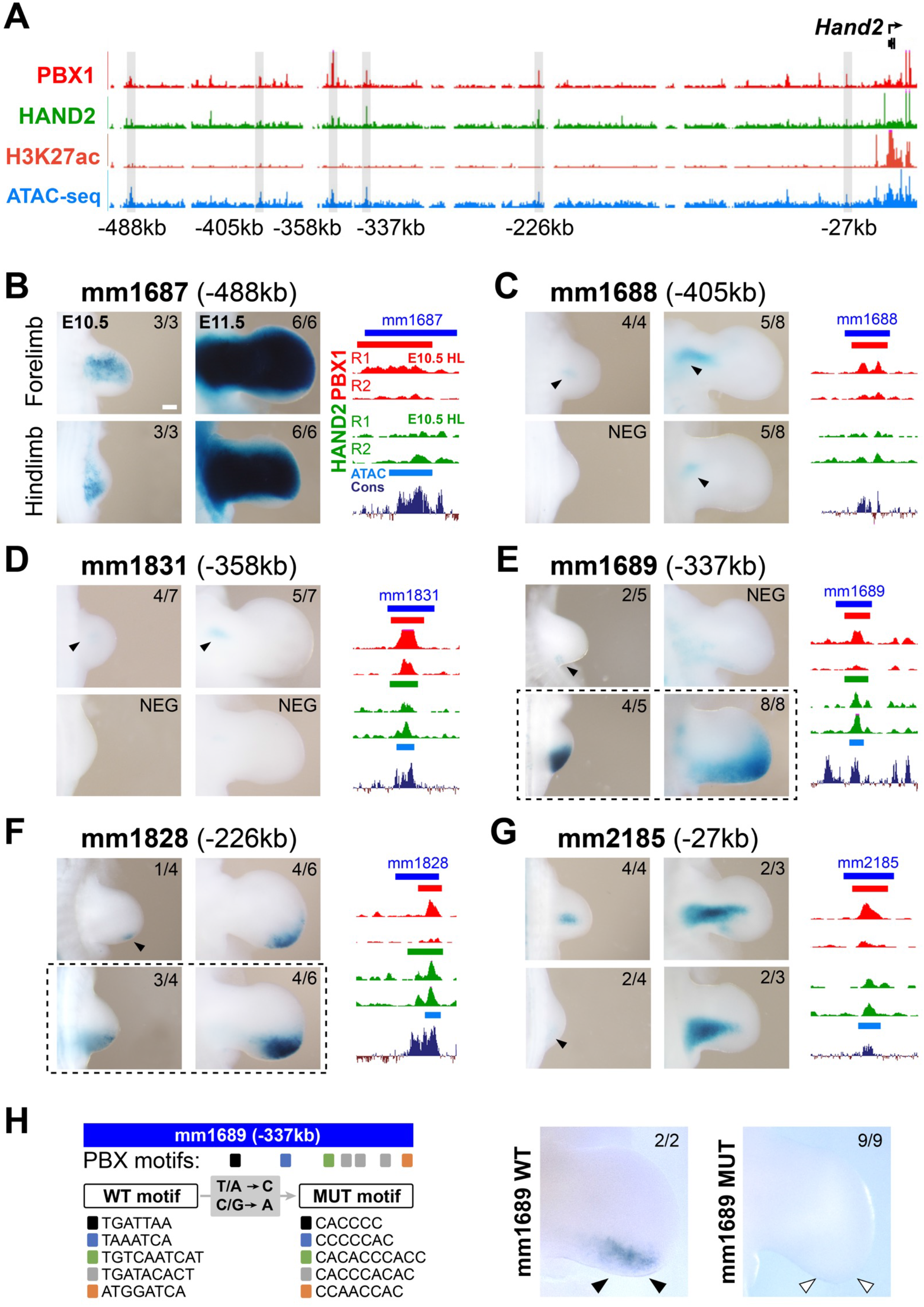
PBX1 directly regulates the enhancer activities of specific CRMs in the *Hand2* TAD. (A) Overview of the genomic signatures of identified candidate limb enhancer elements (grey lines) within the *Hand2* TAD. *Hand2* transcriptional start site (TSS) marked by black arrow (top right). Enhancers listed based on their relative distance to *Hand2* TSS (bottom). (B-G) Analysis of transcriptional enhancer activity in E10.5 and E11.5 forelimb (top panels) and hindlimb (bottom panels) buds using mouse *LacZ* transgenic reporter assays. Left panels: Enhancer activity domains in embryonic forelimbs and hindlimb visualized by *LacZ* staining (blue). Weaker or more restricted domains highlighted by black arrowheads. Numbers at the top right of each panel indicate reproducibility in *LacZ* reporter assay (independent biological replicates with similar results). Right panels: UCSC genome browser tracks depicting replicated PBX1 (red) and HAND2 (green) ChIPseq peaks in E10.5 hindlimb buds for each element tested (blue). ATACseq peak calls (azure) and placental conservation (Cons, dark purple). The 6 enhancers display *bona fide* hindlimb-specific activity in at least one of the two time-points analyzed. Corresponding Vista enhancer IDs (mm: mus musculus) indicated. (H) Analysis of site-directed transcriptional enhancer activity of wild-type (WT) and mutant (MUT) mm1689 enhancer variants in E11.5 hindlimb buds using enSERT. Left Panel: mutagenesis strategy for enhancer mm1689, in which all PBX and PBX-HOX binding sites (WT motif) have been mutagenized (MUT motif). Right Panel: Activity of mm1689 is lost in MUT version. Numbers of embryos with reproducible *LacZ* staining (or lack thereof) in posterior limb bud mesenchyme (arrowheads) over total number of transgenic embryos analyzed (top right). Only transgenic embryos that carried at least two copies of the reporter transgene at the H11 locus were included in our analysis (Kvon et al. 2020) (see also Methods).

### PBX and HAND2 collaboratively control transcriptional regulators of limb patterning

To identify GRNs co-regulated by PBX1/2 and HAND2, we used *Pbx1cKO^Mes^;Pbx2^−/−^* and *Hand2cKO^Mes^* hindlimb buds at E10.5 for bulk RNAseq analysis in comparison to littermate controls (Fig. S7A,B). Statistical analyses identified 1,489 differentially expressed genes (DEGs) in *Pbx1cKO^Mes^*;*Pbx2^−/−^* and 375 DEGs in *Hand2cKO^Mes^* mutant hindlimb buds. Intersection of both transcriptomic datasets, considering only genes that were robustly detected in both sets of experiments, identified 46 DEGs significantly upregulated in hindlimb buds from both mutant mouse lines, defining them as genes repressed by both PBX1/2 and HAND2 (Fig. 5A, left, Fig. S7B and Supplementary Table 4). GO analyses of the PBX1-HAND2 repressed transcriptional targets revealed a significant association with ‘developmental processes’ and ‘transcription factors’ categories (Fig. S7C). In addition, 37 genes were significantly downregulated in both types of mutant hindlimb buds compared to littermate controls (Fig. 5A, left, Fig. S7B,C, and Supplementary Table 4), identifying target genes positively regulated by both PBX1/2 and HAND2 transcription factors. Lastly, 31 DEGs were altered in a discordant manner in the two mutant mouse lines, *i.e.* one transcript was upregulated in *Pbx1cKO^Mes^*;*Pbx2^−/−^* and downregulated in *Hand2cKO^Mes^* mutant hindlimb buds, or *vice versa* (Fig. 5A, left, and Fig. S7B,C). Of note, the GO term ‘transcription factors’ was significantly enriched in all DEG groups, suggesting that PBX1/2 and HAND2 act as master regulators of a downstream GRN comprising transcription factors with critical roles in limb development (Fig. S7C).

**Figure 5.**
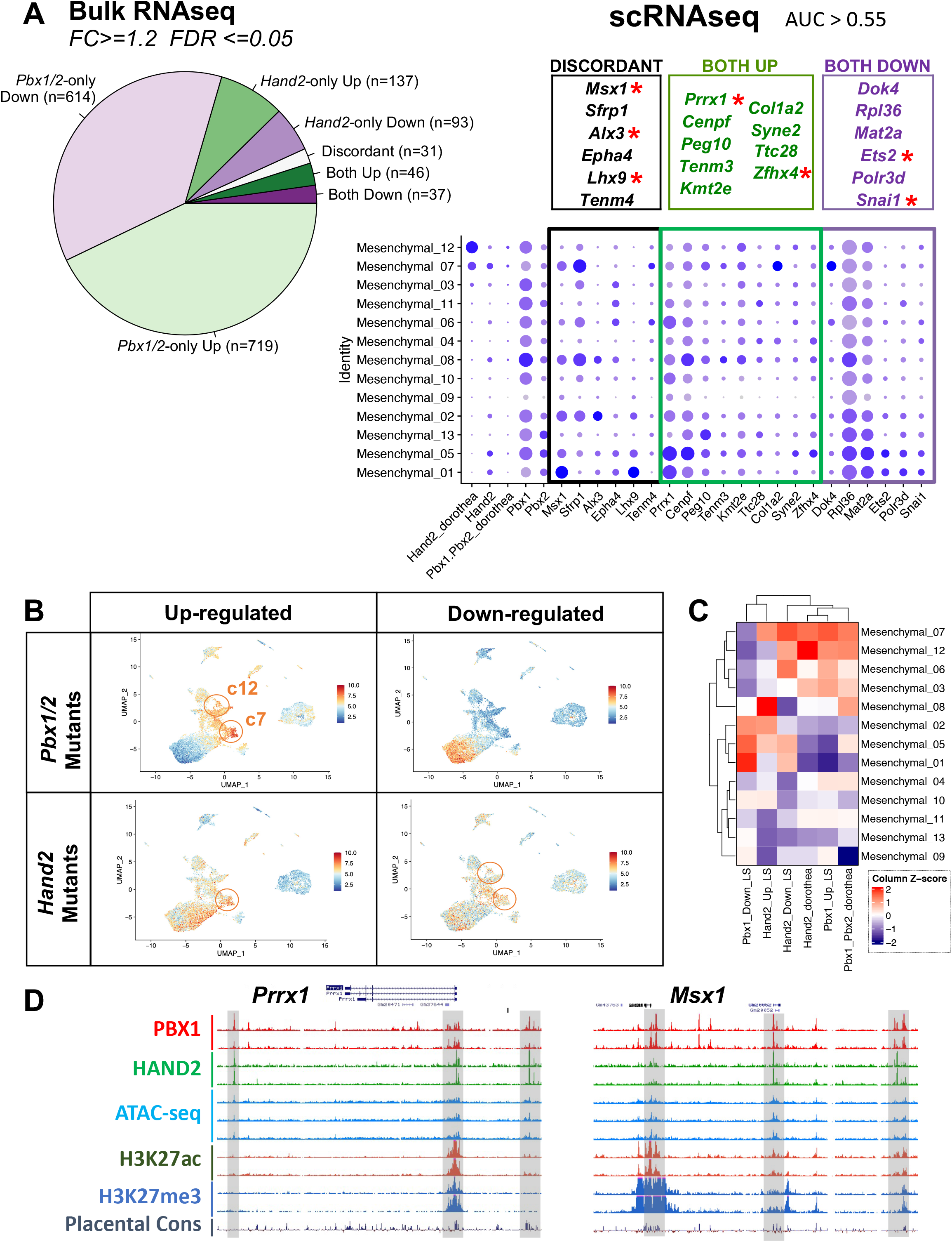
Target genes co-regulated by PBX1/2 and HAND2 are essential for patterning the hindlimb bud. (A) Bulk RNAseq integration of datasets from E10.5 hindlimb buds of *Pbx1cKO^Mes^*;*Pbx2^−/−^* and *Hand2cKO^Mes^* mutants *versus* respective controls. (Left) Pie chart indicating proportion of DEGs in both *Pbx1/2* and *Hand2* mutant hindlimb buds (Both) *versus* littermate controls, or in hindlimb buds from only one of the mutant genotypes (*Pbx1/2*-only and *Hand2*-only) *versus* littermate controls. Number of gene transcripts in each category indicated in brackets. (Right) Intersection of bulk RNAseq with scRNAseq. (Upper Panel) Within boxes, DEGs from both *Pbx1/2*-deficient and *Hand2*-deficient hindlimb buds and also co-expressed in the same cells as *Pbx1/2* and *Hand2* (AUC > 0.55). Transcription factors with known essential roles in limb patterning/morphogenesis highlighted by red asterisk. (Lower Panel) Dot plot of identified target genes showing their expression in the mesenchymal clusters identified by scRNAseq. Dot size represents proportion of cells within a given population that expresses the gene; color intensity indicates average expression level. (B) UMAP in which each cell is color-coded according to the expression of *Pbx1/2 and Hand2* target genes inferred based on RNAseq datasets from *Pbx1/2-* and *Hand2*-mutant hindlimbs. (C) Heatmap showing relative (z-score) median expression of different subsets of *Pbx1/2 and Hand2* target genes in scRNAseq mesenchymal subclusters. (D) UCSC genome browser tracks encompassing *Prrx1* and *Msx1* loci. Tracks: PBX1 and HAND2 ChIPseq, red and green, respectively; ATACseq, aqua; H3K27ac and H3K27me3, brown and blue, respectively; placental conservation, dark purple. Grey rectangles highlight predicted CRMs bound by PBX1 and HAND2.

Upon signal transduction, many transcription factors translocate from the cytoplasm into the nucleus where they interact with other transcription factor complexes and/or bind directly to DNA (Lee and Young 2013). Therefore, we investigated whether the PBX1/2-HAND2 target genes are co-expressed with *Pbx1/2* and *Hand2* in the same hindlimb bud cells by analyzing the mesenchymal clusters in our scRNAseq datasets (based on the statistical threshold ‘area under the curve’ >=0.55 to determine the extent to which each target gene was co-expressed with *Pbx1/2* and/or *Hand2*; see Methods). Remarkably, seven transcription factors with critical functions in limb and/or skeletal development, namely *Msx1* (Bensoussan-Trigano et al. 2011)*, Alx3* (Kuijper et al. 2005)*, Lhx9* (Tzchori et al. 2009)*, Prrx1* (Berge et al. 1998), *Zfhx4* (Nakamura et al. 2021), *Ets2* (Lettice et al. 2012) and *Snai1* (Chen and Gridley 2013; de Frutos et al. 2007) were above this statistical threshold (right panel, red asterisks, Fig. 5A). Genes up-regulated in both *Pbx1/2* and *Hand2* mutants were highly expressed in subcluster 7 (e.g. *Col1a2*), or in subclusters 1 and 5 (e.g. *Prrx1, Msx1*). In contrast, *Ets2, Polr3d*, and *Snai1*, which were downregulated in both mutants, were expressed at higher levels in subclusters 1 and 5. In addition, we used our bulk and sc RNAseq datasets, together with an available computational compendium (Dorothea; Garcia-Alonso et al. 2019; see Methods) to evaluate whether PBX1/2-HAND2 target genes are co-expressed with *Pbx1/2* and *Hand2* in the same cell subpopulations. This analysis detected the largest number of PBX1/2 and HAND2 target genes in cells co-expressing *Pbx1, Pbx2* and *Hand2* within the mesenchymal sub-clusters 7 and 12 (Fig. 5B,C and Fig. S7D). Both promoter and intergenic regions of these target genes interacted with both PBX1 and HAND2 in accessible and active chromatin regions (Fig. 5D). Together, these analyses establish that the co-regulated PBX1/2-HAND2 transcriptional targets required for limb development are: 1) significantly dysregulated in hindlimb buds with mesenchymal-specific loss of *Pbx1/2* and *Hand2;* 2) bound by both PBX1 and HAND2; 3) associated with open and active chromatin; and 4) co-expressed in posterior hindlimb mesenchymal cells at E10.5.

We then examined whether previously defined topological associating domains (TADs, Dixon et al. 2012; Bonev et al. 2017) comprising the identified DEGs exhibited a higher number of putative regulatory elements bound by PBX1 and/or HAND2 than TADs comprising genes whose expression was not altered in *Pbx1cKO^Mes^;Pbx2^−/−^* or *Hand2cKO^Mes^* mutant hindlimbs (Fig. S8). This analysis indicated that distal elements within TADs of DEGs identified in either of the two mutants showed a trend toward higher regulatory scores (based on both the total number and individual strength of the sites contacted by PBX1 and/or HAND2 within the TAD; see Methods) than genes whose expression was not altered in *Pbx1cKO^Mes^;Pbx2^−/−^* and/or *Hand2cKO^Mes^* hindlimb buds. Moreover, the genes within TADs with the highest number and strength of distal regions bound by PBX1 and/or HAND2 were generally upregulated or regulated in a discordant manner in both mutant hindlimb buds (red arrows, Fig. S8). These results indicate that: 1) TADs containing higher numbers of DEGs comprise higher numbers of genomic regions bound by PBX1 and/or HAND2; and 2) genes that are repressed by these two transcriptional regulators are embedded within complex *cis*-regulatory landscapes.

### Genetic interaction between *Pbx1* and *Hand2* is essential for proximo-distal hindlimb and digit patterning

To validate the developmental impact of the novel PBX1/2-HAND2-dependent GRN reconstructed by multi-omics approaches, we assessed its functional importance for normal mouse hindlimb bud patterning and skeletal development by a classical genetic interaction experiment. We generated compound mutant embryos lacking *Pbx1* and one allele of *Hand2* in the hindlimb bud mesenchyme using the *Hoxb6Cre* deleter line (Fig. 6A-G). The hindlimb skeletal morphology was normal in both heterozygous *Pbx1* and *Hand2* embryos (*Pbx1^f/+^;Hoxb6^Cre/+^* and *Hand2^+/−^;Hoxb6^Cre^*) compared to WT controls. In contrast, pelvic malformations and a shorter femur (n=4/4) were detected in *Pbx1^f/f^;Hoxb6^Cre/+^* single homozygous mutant embryos. Compound *Pbx1^f/f^;Hand2^+/−^;Hoxb6^Cre/+^* embryos lacking both copies of *Pbx1* and one copy of *Hand2* in the hindlimb bud mesenchyme revealed a striking genetic interaction in hindlimb skeletal and digit patterning. In particular, *Pbx1^f/f^;Hand2^+/−^;Hoxb6^Cre/+^* hindlimb skeletons exhibited a conspicuously dysmorphic pelvis and shorter femur and severe fibular defects, including agenesis, in combination with variable reduction or loss of posterior digits (Fig. 6D-F’). Despite the different penetrance of the autopod phenotype in compound *Pbx1^f/f^;Hand2^+/−^;Hoxb6^Cre/+^* mutants (Fig. 6D’-F’), no autopod defects were observed in either *Pbx1^f/f^;Hoxb6^Cre/+^* or *Hand2^+/−^;Hoxb6^Cre/+^* hindlimbs. These results establish that genetic interaction of *Pbx1* and *Hand2* is required for normal hindlimb bud patterning.

**Figure 6.**
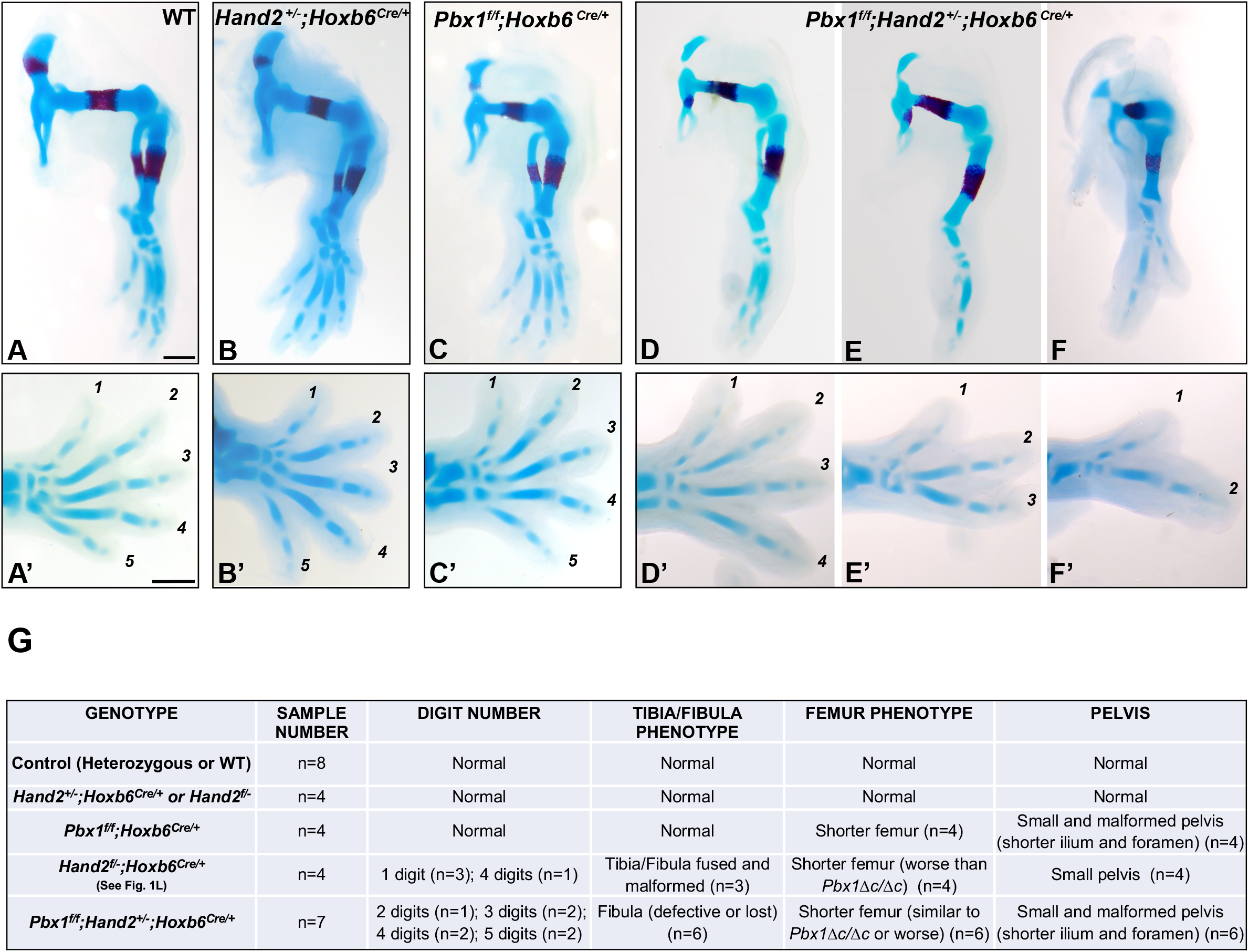
*In vivo* genetic interaction of *Pbx1* and *Hand2* directs patterning of posterior hindlimb skeletal elements. (A-F’) Hindlimb skeletons at E14.5 (blue: cartilage; red: bone). Digits shown from anterior (digit 1) to posterior (digit 5). (A, A’) Representative example of control embryo (Wild-type, WT) (n=8). (B, B’) *Hand2^+/−^;Hoxb6^Cre/+^* hindlimbs with normal morphology. (C, C’) *Pbx1^f/f^; Hoxb6^Cre/+^* hindlimbs with pelvic malformations and shorter femur (n=4/4). (D-F,D’-F’) *Pbx1^f/f^;Hand2^+/−^;Hoxb6^Cre/+^* hindlimbs with malformed pelvis, shorter or truncated femur, defects or loss of fibula, hypoplastic or absent tarsals and metatarsal, and variable loss of posterior digits (n=6/7). Autopod distal-most abnormalities (in tarsal, metatarsal, and phalanges) not observed in either *Pbx1^f/f^;Hoxb6^Cre/+^* or *Hand2^+/−^;Hoxb6^Cre/+^* mutants. Scale bar: 500µm. (G) Table listing total number of embryos analyzed and number of embryos with hindlimb skeletal defects shown in panels A-F.

### HAND2 “selects” a subset of PBX-bound CRMs to confer limb bud patterning specificity

As *Pbx* genes are broadly expressed during embryogenesis and fulfill essential pleiotropic roles during organogenesis (Selleri et al. 2019), we next asked whether context-specific PBX functions are achieved by tissue-specific PBX binding or by cooperativity with different transcription factors that confer tissue-specificity. To this end, we compared PBX genome-wide binding profiles across multiple tissues of the developing embryo. Given the essential roles of PBX proteins in upper lip/primary palate morphogenesis that we reported (Ferretti et al. 2011; Losa et al. 2018), we generated additional PBX1 ChIPseq datasets from the murine midface (MF) at E10.5 and E11.5. In addition, as PBX drives a second branchial arch (BA2)-specific transcriptional program together with MEIS and HOXA2 (Amin et al. 2015), the available PBX1 ChIPseq dataset from E11.5 BA2 was also included in this study. Intersection of the three datasets (PBX1 genome-wide occupancy in the hindlimb bud, MF, and BA2), using only peaks present in both ChIPseq replicates (overlap of at least 1 nt), identified more than 10,000 PBX1-bound genomic regions in each of the tissues analyzed, a large fraction of which was shared across all three embryonic tissues analyzed (Fig. 7A; see Supplementary Table 5 for precise numbers and percentages). The fraction of peaks shared across these datasets is very similar in size to the binding overlap expected across biological replicates of the same genome-wide binding experiment (>50%, Bardet et al. 2011). This points to highly promiscuous PBX binding in different embryonic tissues. GREAT analysis of the PBX binding profiles in hindlimb bud and BA2 revealed significant associations with diverse developmental processes. GO terms associated with limb development (Fig. 7B,C) were included, but they were not the top enriched categories in the hindlimb (or in BA2) dataset. Next, we intersected PBX-bound peaks with HAND2 or HOXA2 binding (Bridoux et al. 2020), and we observed that HAND2 and HOXA2 each occupied only a small subset of PBX peaks. Focusing on the shared HAND2-PBX1 ChIPseq peaks in hindlimb buds revealed that “limb development” and “limb morphogenesis” were among the top three developmental processes (Fig 7D,E). Moreover, HAND2 showed preferential binding to PBX1-bound peaks in the hindlimb bud (48%) compared to BA2 (13%) or MF (18%) tissues (Fig 7D; Supplementary Table 5). In contrast, HOXA2 showed preferential binding to PBX1-bound peaks within BA2 (70%) compared to hindlimb buds (43%) (Fig 7F; Supplementary Table 5). Comparative analysis of HOXA2 binding between BA2 and MF tissues was not conducted, as the MF is a so-called “*Hox*-less” domain (Creuzet et al. 2005). Notably, HAND2- and HOXA2-bound peaks showed only a minimal overlap (12%) and are exclusive (Fig. 7G; Supplementary Table 5). For example, PBX1 binds to similar CRMs in the genomic landscape of *Snail1* and *Zfp503* target genes in hindlimb bud and BA2. However, PBX1 binds the majority of co-bound peaks at both loci with distinct co-factors, HAND2 or HOXA2, in the hindlimb and BA2, respectively (Fig. 7H,I). These data indicate that HAND2 and HOXA2 select for distinct PBX binding events in their domain of expression (hindlimb and BA2, respectively), conferring tissue-specificity to PBX function.

**Figure 7.**
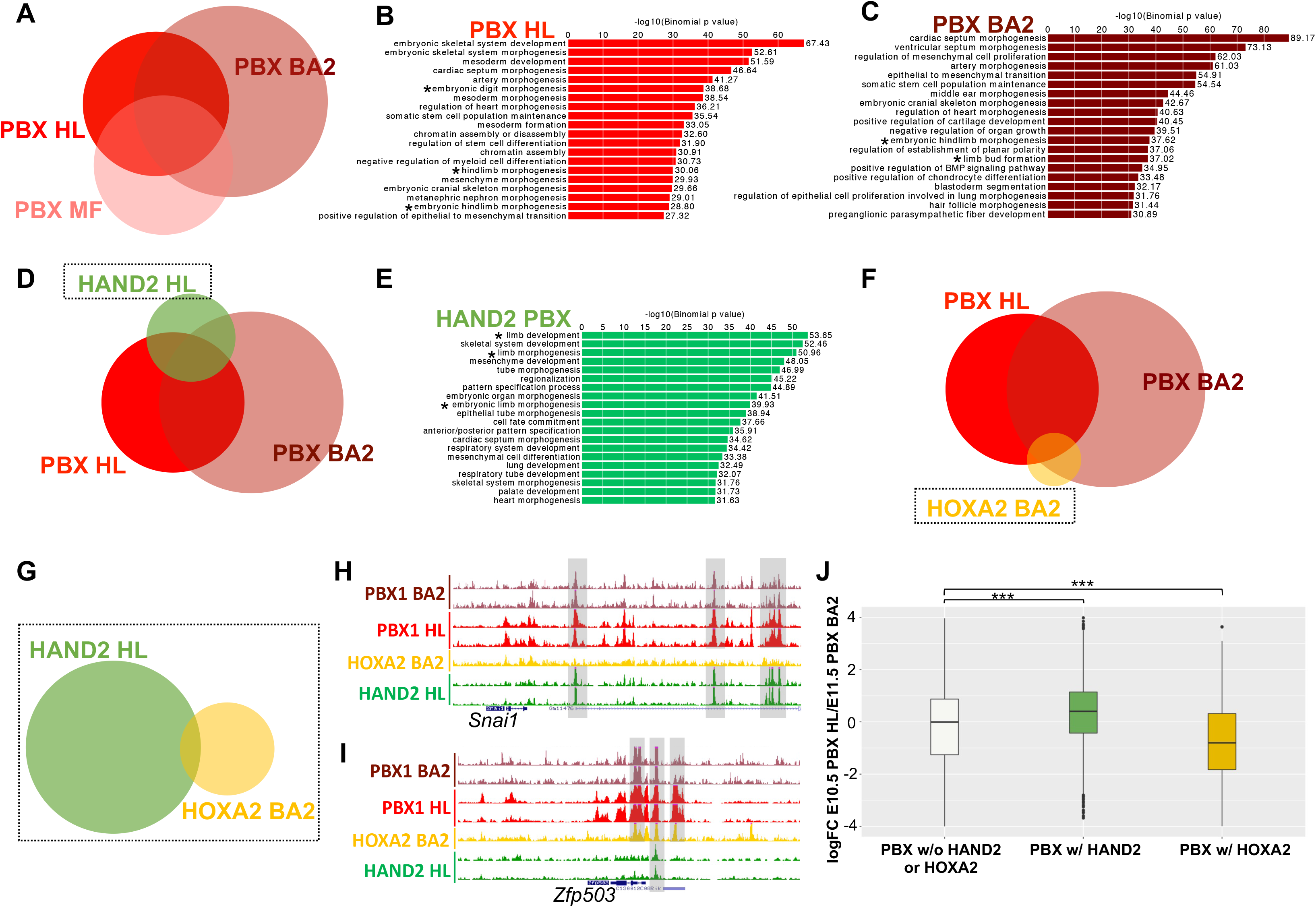
HAND2 selects different subsets of PBX1 peaks to confer early limb patterning functions to PBX1. (A-C) Analysis of genome-wide PBX1 (PBX) binding in E10.5 hindlimb (HL), E11.5 second branchial arch (BA2) and E11.5 midface (MF). Overlap of PBX-bound peaks in hindlimb, BA2 and MF shown in (A). Venn diagram highlighting large overlap between PBX-bound regions across different embryonic tissues (see percentages, Supplementary Table 5). Bar plots showing enriched GO terms associated with genes involved in developmental processes (GREAT) of PBX-bound regions in HL (B) and BA2 (C). Limb development and morphogenesis GO categories (*). Size of bars corresponds to binomial raw (uncorrected) p-values (X axis values). (D) Venn diagram of PBX-bound regions in HL and BA2, with HAND2-bound regions in BA2. (E) Venn diagram of PBX-bound regions in HL and BA2, with HAND2-bound regions in BA2. (F) Overlap of PBX-bound regions in HL and BA2, with HOXA2-bound regions in BA2. (G) Venn diagram of regions bound by HOXA2 in BA2 with those bound by HAND2 in HL. (H,I) UCSC genome browser tracks (mm10) showing the generated ChIPseq datasets from PBX1 BA2 (brown); PBX1 HL (red); HOXA2 BA2 (yellow); and HAND2 HL (green) at the *Snai1* (H) and *Zfp503* (I) loci. Shared peaks highlighted by grey rectangles. (J) Boxplots of ratio (log2Fold Change; logFC) of normalized PBX1 ChIPseq signals between HL and BA2 at indicated subsets of PBX-bound regions.

Lastly, comparing the hindlimb bud to BA2 tissue, we quantified PBX1 binding enrichment of peaks that also overlap with HAND2 or HOXA2 binding. We found that PBX1-HAND2 co-bound peaks in the hindlimb bud are significantly more enriched relative to PBX1 peaks not overlapping HAND2 or HOXA2 binding (p < 2.2e-16) (Fig. 7J). Similarly, co-binding with HOXA2 significantly increased PBX1 binding in BA2 (p < 2.2e-16) (Fig. 7J). In summary, these results demonstrate that promiscuous PBX transcription factors occupy a vast pool of shared genomic regions during embryonic development, but acquire restrained tissue specificity via cooperation with distinct transcriptional regulators, such as HAND2 in the developing hindlimb.

## DISCUSSION

Disturbance of the gene expression programs that specify cell fates and control tissue patterning, as well as morphogenesis of organs, causes congenital malformations. As transcription factors rarely work in isolation, the regulation of gene expression is often mediated by collaborative interactions of transcription factor within complexes (Morgunova and Taipale 2017). Despite decades of research, it remains unclear how spatio-temporal interactions among transcription factors and select cofactors achieve functional specificity and how transcription factor complexes regulate their target genes through multiple CRMs such as enhancers and repressors. Our previous studies reported that, despite broad expression of *Pbx* genes during embryonic development, their encoded transcription factors control distinct target and effector genes in different tissues and organs, including the limb bud (Selleri et al. 2019; Capellini et al. 2011b; Hurtado et al. 2015; Brendolan et al. 2005; Golonzhka et al. 2015; McCulley et al. 2018; Stankunas et al. 2008; Welsh et al. 2018). Our research highlighted the essential roles of *Pbx1/2* during limb bud development (Capellini et al. 2011b; Selleri et al. 2019), but did not identify the cell type(s) that require PBX function or the time window during which limb bud development depends on PBX. Beyond the phenotypic characterization and identification of target genes, our previous studies did not provide insight into how PBX factors attain functional specificity during development despite their widespread localization in the embryo. The genome-wide and single-cell experimental strategies employed allowed us to address these critical issues and provided new insight into broader questions related to transcription factor promiscuity and specificity. We previously reported that inactivation of both *Pbx1* and *Pbx2* results in more severe phenotypes in mouse hindlimbs than forelimbs (Capellini et al. 2008, 2006), likely as a consequence of functional redundancy among *Pbx* family members, as *Pbx3* is not expressed in hindlimbs. Thus, in this study we chose the mouse hindlimb bud to gain a mechanistic understanding of how TALE PBX homeoproteins control specific transcriptional programs *in vivo* in a context-dependent manner.

The combination of tissue and temporally-controlled gene inactivation in the mouse with multi-omics approaches using dissected hindlimb buds established that *Pbx1/2* are dispensable for AER formation and function, but essential in the mesenchyme for normal hindlimb bud initiation and early patterning. Thereafter, PBX1/2 become dispensable in the hindlimb mesenchyme, pointing to their crucial roles in mesodermal progenitors during the onset of limb bud development. Consistent with essential PBX1/2 functions in mesoderm patterning, we reported that they are required to control *Polycomb* and *Hox* gene expression in the paraxial mesoderm, which gives rise to the axial skeleton (Capellini et al. 2008). Of note, a recent study showed that induction of the paraxial mesoderm relies on a BRACHYURY-TALE-HOX gene code (Mariani et al. 2021). In this context, TALE-HOX transcription factors establish a chromatin landscape permissive for recruitment of the WNT-effector LEF1, which in turn unlocks WNT-mediated transcriptional programs that drive paraxial mesodermal fates. Similarly, our study establishes that PBX1/2 homeoproteins, in cooperation with tissue-specific cofactors, execute essential roles in the specification and patterning of mesodermal lineages in both axial and limb bud mesoderm.

Limb bud mesenchyme-specific deletion of either *Pbx1/2* or *Hand2* causes similar limb skeletal defects and comparable cellular and molecular alterations. In addition, the phenotypes of mouse embryos lacking either *Pbx1/2* or *Hand2* in the limb mesenchyme phenocopy the morphological and molecular defects of mouse embryos lacking *Shh* (Chiang et al. 2001) or its limb ZRS enhancer (Sagai et al. 2005), resulting in loss of posterior digits. We and others reported that activation of *Shh* localized expression in the posterior limb bud mesoderm is regulated by interactions of transcriptional regulators, including HOX, PBX, HAND2, and ETS with the ZRS, whereas TWIST1, ETV, and GATA factors prevent anterior ectopic activation of *Shh* expression (Galli et al. 2010; Kmita et al. 2005; Kozhemyakina et al. 2014; Lettice et al. 2012, 2014; Mao et al. 2009; Capellini et al. 2006). Here, we demonstrate that *Pbx1/2* and *Hand2* cooperate to regulate a GRN that controls the initiation of hindlimb bud patterning. Lastly, we establish that PBX1 regulates *Hand2* expression via the control of select CRMs active within the posterior hindlimb bud mesenchyme. Indeed, mutations of all PBX and PBX-HOX binding sites for the mm1689 CRM demonstrated that PBX is essential for its activity. Of note, it was recently reported that deletion of this enhancer, which comprises also MEIS and HOXD13 binding sites, results in loss of *Hand2* expression in murine limb buds (Delgado et al. 2021), strongly arguing that PBX is part of the transcriptional complex that is required for *Hand2* activation. Together, these findings provide the first mechanistic evidence that explains how the loss of *Pbx1/2* or *Hand2* results in similar skeletal, cellular, and molecular alterations during limb bud development.

Transcription factors that act within the same GRN must be co-expressed within the same cells, as they translocate into the nucleus to regulate common downstream target gene expression (Lee and Young 2013). A recent study based on synthetic GRN modelling and randomization tests reported that widespread co-expression emerges in regulatory networks and is a direct indicator of active co-regulation within a given cellular context (Yin et al. 2021). Combination of scRNASeq and immunofluorescence on hindlimb buds provides compelling evidence that both *Pbx1/2* and *Hand2* transcripts are co-expressed within restricted mesenchymal subpopulations in the posterior-proximal hindlimb bud, where PBX1 and HAND2 proteins also co-localize in nuclei. The PBX1/2-HAND2-dependent GRN built here comprises target genes that encode critical limb developmental regulators within well-defined epigenetic landscapes. To summarize, this spatio-temporally constrained GRN achieves superior organismal- and tissue-level resolution within a confined hindlimb bud domain (Fig. 8).

**Figure 8.**
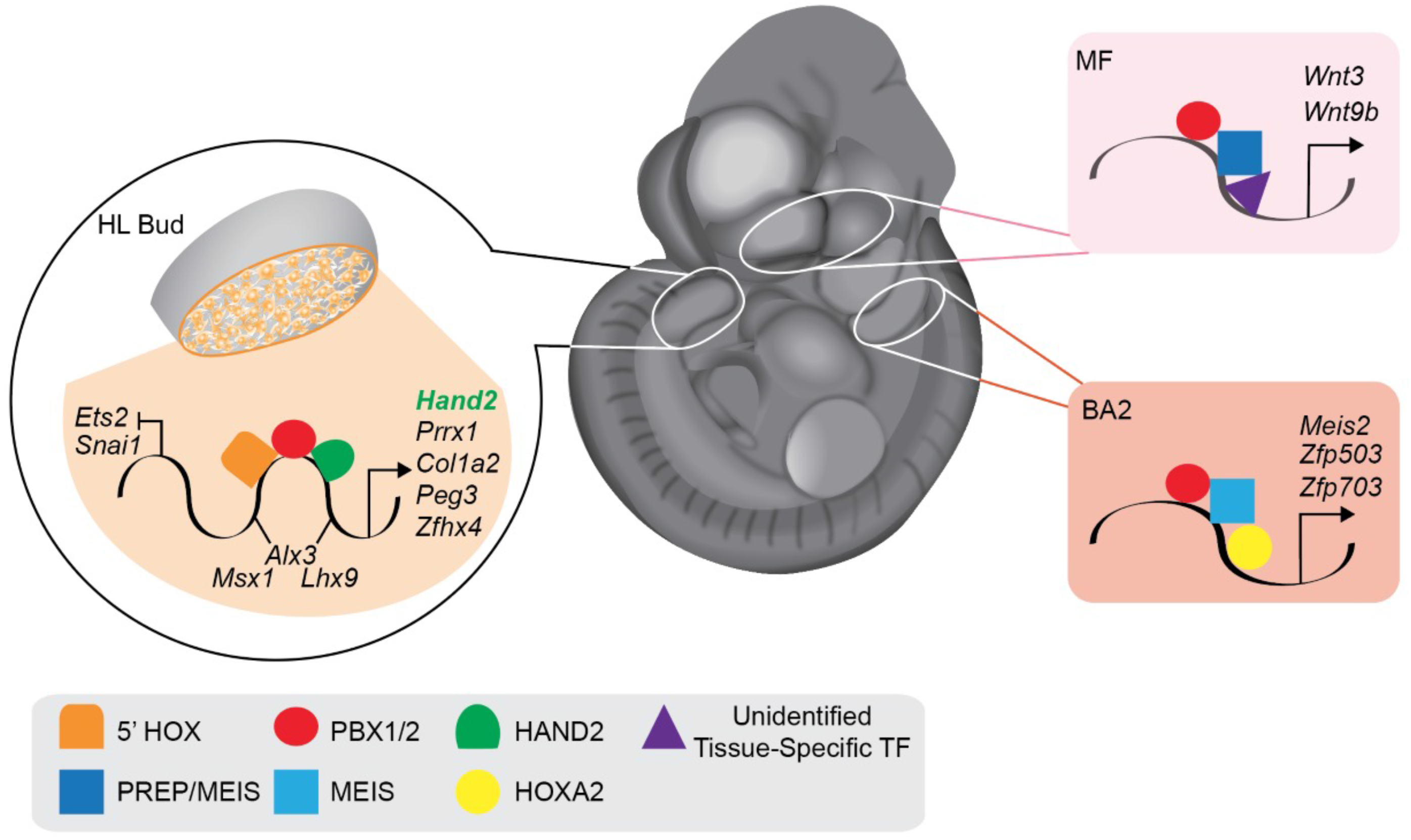
Promiscuous genome-wide PBX binding acquires tissue-specific developmental functions via interactions with distinct cofactors. Model depicting how PBX homeoproteins must rely on DNA binding partnerships with different cofactors to execute context-dependent developmental functions. In the midface (MF), a “*Hox*-less” domain, PBX1/2 together with PREP/MEIS bind regulatory elements of *Wnt3* and *Wnt9b* (pink-shaded area) to direct their expression during upper lip/primary palate fusion (Ferretti et al. 2011). In the second branchial arch (BA2), PBX1/2 activate a BA2-specific transcriptional program by binding the *Zfp503* and *Zfp703* loci with HOXA2 and MEIS (ochre-shaded area) (Amin et al. 2015). In restricted subset of posterior hindlimb bud mesenchymal cells (orange shading reflects co-expression of *Pbx1/2*, red, and *Hand2,* green), a PBX1/2-directed GRN acquires early limb patterning functions via interaction with HAND2. This spatio-temporally constrained GRN (orange-shaded area) activates known limb transcriptional regulators, such as *Prrx1* and *Hand2* itself, and represses other limb transcription factors, such as *Snai1* and *Ets2*. In contrast, PBX1/2 and HAND2 regulate other target genes (*Alx3, Msx1, Lhx9*) in a discordant manner.

It was recently reported that *Meis1/2* genes, which are broadly expressed in the mouse embryo and encode TALE homeodomain proteins that can dimerize with PBX (Moens and Selleri 2006; Selleri et al. 2019), have also essential roles during early limb bud development. Their genetic inactivation in the mouse results in distal abnormalities including loss of posterior digits (Delgado et al. 2021, 2020), similar to *Pbx1/2*-deficient mice. Furthermore, it was shown that MEIS and TBX transcription factors control a limb-specific GRN by co-regulating enhancers associated with genes essential for limb bud initiation such as *Fgf10* (Delgado et al. 2021). While this study showed that expression of *Hand2* and *Shh* is altered in *Meis1/2*-deficient hindlimbs, it remains to be determined whether MEIS proteins are part of the same PBX1/2-HAND2-directed GRN, or of a larger shared GRN. Our *de novo* motif analysis provides evidence that PBX1 can bind DNA without HAND2, but likely together with other TALE factors such as MEIS1/PREP1 in hindlimb buds (see Fig. 3H, third column), in agreement with ChIPseq experiments conducted on whole embryos (Penkov et al. 2013). Therefore, we envisage a scenario whereby, together with PBX1/2 and HAND2, MEIS proteins are part of the same transcriptional complex with other homeodomain transcription factors (see Fig. S4H) that participate in the regulation of shared target genes with essential functions in hindlimb bud patterning. Supporting this view, it was reported that MEIS and PBX largely occupy the same genomic regions in the branchial arches (Bridoux et al. 2020; Amin et al. 2015). Of note, transcription factor cooperativity does not only rely on direct protein-protein interaction but can also be indirect. In this case, the transcription factors do not interact but stabilize each other by binding to the same genomic regions, thus competing with nucleosomes (Mirny 2010; Moyle-Heyrman et al. 2011; Morgunova and Taipale 2017).

Several studies have attributed ‘pioneer factor’ functions to PBX proteins (Choe et al. 2014, 2009; Grebbin et al. 2016; Grebbin and Schulte 2017). For example, it was proposed that during early development PBX may penetrate repressive chromatin to mark specific genes for transcriptional activation as part of multimeric complexes (Berkes et al. 2004; Sagerström 2004). Furthermore, it was shown that during early zebrafish embryogenesis TALE regulators access promoters, thereby facilitating chromatin accessibility of transcriptionally inactive genes, and subsequently HOX proteins are required to initiate transcription (Choe et al. 2014). In the mouse, it was reported that during neural differentiation PBX1 binds the promoter/proximal enhancer of doublecortin (*Dcx*) in undifferentiated neural progenitors, when chromatin is still compacted by histone H1 and long before *Dcx* is expressed (Grebbin et al. 2016). Our study generates new genome-wide evidence in support of the putative pioneer factor functions of PBX proteins: 1) PBX1 without HAND2 binds to regions that are significantly more accessible and more enriched with H3K27ac than regions bound by HAND2 without PBX1 (see Fig. 3I); 2) there is a significant association between repressed promoter elements marked by H3K27me3 and PBX1 binding without HAND2 (see Fig. S5D); and 3) in different embryonic tissues, including hindlimb bud, MF, and BA2, DNA regions bound by PBX1 are largely overlapping (see Fig. 7). Nonetheless, in order to unequivocally assign pioneer factor roles to PBX proteins it will be critical to establish that PBX-marked genes are not already primed for transcriptional activation by pre-existing histone modifications (Iwafuchi-Doi and Zaret 2014; Donaghey et al. 2018) prior to PBX binding. As PBX1/2 are required in the limb mesenchyme during hindlimb bud initiation, our study supports the notion that these TALE regulators have critical functions in multipotent progenitor cell populations (Mariani et al. 2021). Of note, while in the mouse embryo the onset of *Hox* gene expression occurs only during gastrulation (Deschamps and Duboule 2017), *Pbx* genes are expressed earlier, namely in the blastocyst and morula onward (Sonnet et al. 2012). Furthermore, *Pbx* gene transcripts are already detected in both mouse and human embryonic stem cells (Chan et al. 2009; Jürgens et al. 2009), consistent with putative pioneer factor functions.

PBX homeodomain proteins of the TALE superclass (Moens and Selleri 2006) are characterized by the insertion of a three-amino-acid loop in the homeodomain, a highly conserved DNA binding moiety shared by hundreds of transcription factors (Bürglin and Affolter 2015; Bobola and Merabet 2017), which typically is not sufficient to direct transcription factors to their functional targets. Our study shows that PBX proteins bind indiscriminately to a vast number of CRMs shared across diverse embryonic tissues. Thus, binding of PBX TALE proteins alone seems insufficient for context-dependent functional activity, whereas PBX1 binding acquires tissue-specific functions via combinatorial interactions with different transcription factors, such as HAND2, which confers limb bud specificity (see Fig. 8). A similar functional model defining how TALE factors can activate target genes in different contexts was proposed for MEIS (Bridoux et al. 2020; Amin et al. 2015). Based on our findings (see Fig. 3 and Fig. 7), one can envisage a scenario whereby increased accessibility reflects higher DNA binding affinity of PBX in the presence of HAND2, and ultimately longer residence time on chromatin. Of note, the duration of transcription factor binding to DNA -or dwell time-positively correlates with downstream transcriptional output (Donovan et al. 2019; Lickwar et al. 2012).

PBX proteins have long been considered to serve primarily as cofactors for HOX proteins, and their heterodimerization with PBX has been classically proposed as a mechanism by which HOX proteins acquire DNA-binding selectivity and specificity (Pöpperl et al. 1995; de Kumar et al. 2017; Parker et al. 2019; Ryoo and Mann 1999; Mann et al. 2009). However, it always remained difficult to envisage how transcription factors such as PBX homeoproteins, encoded by genes with widespread expression in the embryo, can confer functional specificity to HOX proteins, that display domain-restricted localization (Selleri et al. 2019). Challenging this widely accepted model, we reported that PBX transcriptional regulators can hierarchically control *Hox* gene expression in limb buds (Capellini et al. 2006) and also function in “*Hox*-less” embryonic domains, such as the developing head (Losa et al. 2018; Welsh et al. 2018; Ferretti et al. 2011). Using the limb bud as a model system, our study addresses the critical lingering question of how PBX proteins achieve developmental specificity. Our research provides novel evidence that interaction with select cofactors, such as HAND2 in hindlimb buds, restrains promiscuous PBX proteins directing them to execute specific developmental functions in distinct embryonic tissues.

## MATERIALS AND METHODS

### Mice

Mutant alleles used in this study were previously described and the conditions for genotyping were reported: conditional *Pbx1* (Koss et al. 2012)*, Pbx2* (Selleri et al. 2004)*, Hand2^3xFLAG^* allele (Osterwalder et al. 2014), constitutive and conditional *Hand2* (Galli et al. 2010), *Hoxb6Cre* (Lowe et al. 2000), *Hoxb6CRE-ERT* (Nguyen et al. 2009)*, Msx2Cre* (Liu et al. 1994; Sun et al. 2000). Experiments on *Pbx1*/*Pbx2* mice were performed following Weill Cornell Medical College and UCSF IACUC guidelines and experimental procedures concerning housing, husbandry, and welfare. Animal studies conducted in Switzerland involving *Hand2* and *Pbx*/*Hand2* compound mutant mice and embryos were approved by the Regional Commission on Animal Experimentation and the Cantonal Veterinary Office of the city of Basel. Animal work at Lawrence Berkeley National Laboratory (LBNL) was reviewed and approved by the LBNL Animal Welfare Committee. Stages of all embryos analyzed are indicated in all figures and figure legends. Wild-type (Swiss Webster) mice were purchased from Charles River Laboratories and were time-mated to obtain embryos for microdissections. Mouse embryos were collected from pregnant females via cesarean section at the described timepoint following observation of a vaginal plug. Noon the day of the plug was considered E0.5.

### Skeletal preparations

Skeletal preparations were performed as previously described (Capellini et al. 2006, 2010; Galli et al. 2010).

### Whole-mount ISH

Whole-mount *in situ* hybridization was performed on E11.5 embryos as previously described (Capellini et al. 2008, 2006; Galli et al. 2010). At least three embryos for each genotype were analyzed by whole-mount *in situ* hybridization.

### Immunofluorescence

Embryos were harvested at specified stages, fixed overnight at 4°C in 4% PFA in PBS, rinsed twice in PBS, and cryoprotected in 30% sucrose overnight at 4°C. Subsequently, they were embedded in OCT compound and cryosectioned at 12μm per section. Slides were blocked for 1h with 10% fetal bovine serum (FBS)/PBS and incubated overnight in 0.1% bovine serum albumin (BSA) with primary antibodies (Ab). Primary Abs used for immunofluorescence were: PBX1 (Cell Signalling, #4342, 1:200), FLAG M2 (Sigma, F1804, 1:500). Primary Ab binding was detected by AlexaFluor-conjugated econdary Abs (Invitrogen, 1:400-1:1000). Nuclei were stained with DAPI (Sigma). Fluorescence imaging was performed using a Leica SP5 confocal microscope.

### scRNAseq

Microdissected hindlimb tissues were collected from 10 embryos at E10.5 (37-40 somites) in cold PBS. Tissues were dissociated to single cells using an enzymatic cocktail of Liberase and DNase I for 10 minutes at 37 degrees. Cells were passed through a 45 µm strainer to remove clumps and ensure a single cell suspension. Dead cells were eliminated with a ‘Dead Cell Removal Kit’ with magnetic beads (MACS, Milteny Biotech). A single cell suspension of pooled live hindlimb cells were loaded into one well for single-cell capture using the Chromium Single Cell 3’ Reagent Kit V2 (10X Genomics). Library preparation for each sample was also performed using the Chromium Single Cell 3’ Reagent Kit V2, and each sample was given a unique i7 index. Libraries were pooled and subjected to sequencing in an Illumina NovaSeq sequencer. Additional details in Suppl. Methods.

### Bulk RNAseq

Hindlimb buds from mutant and littermate controls were dissected individually at the indicated gestational days and snap frozen in dry ice or liquid nitrogen. After genotyping, each biological replicate consisted of hindlimb buds from one individual mutant or littermate embryo, with a total of 3 biological replicates per genotype. RNA was extracted using the RNeasy Plus Micro kit (Qiagen, #74034) and RNA quantification performed using the Qubit RNA HS Assay Kit (Invitrogen, #Q32852). Quality control of input RNA was performed using the RNA 6000 Pico kit (Agilent, #5067–1513) on a 2100 Bioanalyzer (Agilent) or the Fragment Analyzer (Advanced Analytical) High Sensitivity RNA kit. All RNA samples for library preparation had RIN>9. RNA sequencing libraries were prepared from 100 ng of input RNA using the non-directional kit NEBNext Ultra™ II RNA Library Prep Kit for Illumina (NEB, #E7775) with the NEBNext Poly(A) mRNA Magnetic Isolation Module (NEB, #E7490) to capture polyA RNAs. Library size and quality was checked using an Agilent 2100 Bioanalyzer with the High Sensitivity DNA kit (Agilent, #5067–4626) or the Fragment Analyzer CRISPR discovery kit. Concentration of the libraries was determined with the QuBit dsDNA HS Assay kit (Invitrogen, #Q32854). *Pbx1/2* libraries were sequenced in an Illumina HiSeq 4000 to generate 50 base pair single-end reads. *Hand2* libraries were sequenced using NextSeq500 to generate 75 single-end reads. Additional details in Suppl. Methods.

### ChIPseq

Embryonic hindlimb buds and embryonic midfaces (the latter dissected as described in Losa et al. 2018) were isolated from wild-type Swiss Webster E10.5 and E11.5 mouse embryos, and immediately crosslinked for 10 min in 1% formaldehyde (Electron Microscopy Sciences, #15710). ChIP assays were performed as reported (Amin et al. 2015; Losa et al. 2017, 2018). The crosslinked material was sonicated to 200-500 bp DNA fragments with a Diagenode Bioruptor or Covaris S220 sonicator. Each ChIPseq assay was performed from pooling 60-80 embryonic hindlimbs, by O/N incubation with specific antibodies (5 µg) at 4°C followed by 30 min incubation with Dynabeads protein A (PBX1, H3K27ac and H3K27me3 ChIPseq) or 60 min incubation with Dynabeads protein G (HAND2^3XF^ ChIPseq). IP and input DNA were purified using the MicroChIP DiaPure kit (Diagenode, #C03040001). Antibodies used were: PBX1 (Cell Signaling, #4243S), FLAG for HAND2 (Sigma, F1804), H3K27ac (Abcam, ab4729) and H3K27me3 (Millipore, #07-449). Each chromatin immunoprecipitation assay was conducted in two independent biological replicates. Following ChIP, DNA libraries were constructed using the MicroPlex Library Preparation Kit v2 (Diagenode, C05010012). Libraries were sequenced in an Illumina HiSeq 4000 to generate 50 base pair single-end reads. Additional details in Suppl. Methods.

### ATACseq

We used the Assay for Transposase Accessible Chromatin (ATACseq) protocol, as originally described (Buenrostro et al. 2013) with minor modifications. About 75 000 single cells from a pair of mouse hindlimb buds were used and n=3 biological replicates were analyzed. ATACseq reads were processed as described for ChIPseq. Peak calling was also performed with MACS v1.4, using the following parameters: --gsize=mm --bw=150 --nomodel --nolambda --shiftsize=75. Evidence from the three replicates was combined using the same approach described for ChIPseq. ATACseq reads were processed as described for ChIPseq. Peak calling was also performed with MACS v1.4, using the following parameters: --gsize=mm --bw=150 --nomodel --nolambda --shiftsize=75. Evidence from the three replicates was combined using the approach described for ChIPseq.

### *Hand2* candidate enhancer identification and selection for analysis of *LacZ* activity

We intersected our ChIPseq and ATACseq datasets from E10.5 hindlimb buds to identify all regulatory elements within the *Hand2* TAD (Bonev et al. 2017). We selected all PBX1-bound replicated peaks showing at least a 15-fold-enrichment, excluding promoter regions. To identify the complement of transcriptional enhancers controlling *Hand2* expression in the limb mesenchyme, we intersected evolutionarily conserved elements with PBX1-interacting regions enriched for H3K27ac marks and ATACseq peaks.

### Transgenic mouse reporter assays

Transgenic mouse assays at LBNL were performed in *Mus musculus* FVB strain mice. Animals of both sexes were used in these analyses. Sample size selection and randomization strategies were conducted as follows: sample sizes were selected empirically based on previous experience in transgenic mouse assays for >3,000 total putative enhancers (VISTA Enhancer Browser: https://enhancer.lbl.gov/). Mouse embryos were excluded from further analysis if they did not encode the reporter transgene or if the developmental stage was not correct. All transgenic mice were treated with identical experimental conditions. Randomization and experimenter blinding were unnecessary and not performed. For validation of *in vivo* limb enhancer activities conventional transgenic mouse *LacZ* reporter assays involving an hsp68 minimal promoter (*Hsp68*-*LacZ*) were performed as previously described (Kothary et al. 1989; Osterwalder et al. 2022). For comparison of wildtype and mutagenized enhancer versions (mm1689) enSERT was used for site-directed insertion of transgenic constructs at the H11 safe-harbor locus (Kvon et al. 2020). Hereby, Cas9 and sgRNAs were co-injected into the pronucleus of FVB single cell-stage mouse embryos (E0.5) with the targeting vector encoding a candidate enhancer element upstream of the *Shh*-promoter-*LacZ* reporter cassette (*Shh-LacZ*) (Kvon et al. 2020). Related genomic enhancer coordinates are listed in Supplementary Table 3. Predicted enhancer elements were PCR-amplified from mouse genomic DNA (Clontech) and cloned into the respective *LacZ* expression vector (Osterwalder et al. 2022). Embryos were excluded from further analysis if they did not contain a reporter transgene in tandem. CD-1 females served as pseudo-pregnant recipients for embryo transfer to produce transgenic embryos which were collected at E10.5 or E11.5 and stained with X-gal using standard techniques (Osterwalder et al. 2022).

## Data availability

ChIPseq, ATACseq, scRNAseq and RNAseq datasets have been deposited in the NCBI GEO database under the identifier GSE197859. Additional data supporting the findings of this study are available from the corresponding author upon request.

## Acknowledgements

We are grateful to Mr. R. Aho for all artwork; Drs. D. Penkov and F. Blasi for input on ChIPseq protocols and hospitality in their laboratory; A. Baur and O. Romashkina for expert technical assistance; and A. Offinger and her team for mouse care. We thank members of the Selleri laboratory and Drs. G. Panagiotakos, J. Bush, R. Blelloch, D. Lim, Y. Shen, and D. Wagner for useful discussions. Work funded by the National Institutes of Health (grants R01HD043997 to LS; R01DE028599, UM1HG009421, and R01HG003988 to AV); by the Swiss National Science Foundation grant no. 310030_184734 to RZ and AZ and grant PCEFP3_186993 to MO; by the University of California, San Francisco Chancellor’s Funds to LS; and by core funding from the University of Basel to RZ and AZ. Research at the E.O. Lawrence Berkeley National Laboratory performed under US Department of Energy contract DE-AC02-05CH11231, University of California. ML was recipient of postdoctoral fellowship from the American Association for Anatomy and IB was funded by Imperial College Research Fellowship and by Medical University of Vienna.

## Author contributions

LS and RZ designed the study; AZ designed the genetic interaction experiment; ML, MO, JDB, BC, AG, AM and TC designed and performed experiments under the supervision of LS, RZ, AZ, AV, DD and NB; IB, PZ, and NB performed bioinformatic analysis of all data sets. JZ and SM provided an unpublished mouse line and intellectual input. Based on the draft written by ML and LS, the manuscript was critically reviewed by RZ, AZ, IB, MO, TC, SM, NB and AV. All authors read the final version of the manuscript.

## REFERENCES

Amândio AR, Beccari L, Lopez-Delisle L, Mascrez B, Zakany J, Gitto S, Duboule D. 2021. Sequential in cis mutagenesis in vivo reveals various functions for CTCF sites at the mouse HoxD cluster. Genes and Development 35: 1490–1509. http://genesdev.cshlp.org/content/35/21-22/1490.full.

Amin S, Donaldson IJ, Zannino DA, Hensman J, Rattray M, Losa M, Spitz F, Ladam F, Sagerström C, Bobola N. 2015. Hoxa2 Selectively Enhances Meis Binding to Change a Branchial Arch Ground State. Developmental Cell 32: 265–277. http://www.cell.com/article/S1534580714008454/fulltext.

Bardet AF, He Q, Zeitlinger J, Stark A. 2011. A computational pipeline for comparative ChIP-seq analyses. Nature Protocols 2012 7:1 7: 45–61. https://www.nature.com/articles/nprot.2011.420.

Bensoussan-Trigano V, Lallemand Y, saint Cloment C, Robert B. 2011. Msx1 and Msx2 in limb mesenchyme modulate digit number and identity. Developmental Dynamics 240: 1190–1202. https://pubmed.ncbi.nlm.nih.gov/21465616/.

Berge D ten, Brouwer A, Korving J, Martin JF, Meijlink F. 1998. Prx1 and Prx2 in skeletogenesis: roles in the craniofacial region, inner ear and limbs. Development 125: 3831–3842.

Berkes CA, Bergstrom DA, Penn BH, Seaver KJ, Knoepfler PS, Tapscott SJ. 2004. Pbx marks genes for activation by MyoD indicating a role for a homeodomain protein in establishing myogenic potential. Molecular cell 14: 465–477. https://pubmed.ncbi.nlm.nih.gov/15149596/.

Bobola N, Merabet S. 2017. Homeodomain proteins in action: similar DNA binding preferences, highly variable connectivity. Current opinion in genetics & development 43: 1–8. https://pubmed.ncbi.nlm.nih.gov/27768937/

Bolt CC, Duboule D. 2020. The regulatory landscapes of developmental genes. Development 147.

Bonev B, Cohen NM, Szabo Q, Fritsch L, Papadopoulos GL, Lubling Y, Xu X, Lv X, Hugnot J-P, Tanay A, et al. 2017. Multiscale 3D Genome Rewiring during Mouse Neural Development. Cell 171: 557. https://pmc/articles/PMC5651218/.

Brendolan A, Ferretti E, Salsi V, Moses K, Quaggin S, Blasi F, Cleary ML, Selleri L. 2005. A Pbx1-dependent genetic and transcriptional network regulates spleen ontogeny. Development 132: 3113–3126. https://pubmed.ncbi.nlm.nih.gov/15944191/.

Bridoux L, Zarrineh P, Mallen J, Phuycharoen M, Latorre V, Ladam F, Losa M, Baker SM, Sagerstrom C, Mace KA, et al. 2020. HOX paralogs selectively convert binding of ubiquitous transcription factors into tissue-specific patterns of enhancer activation. PLOS Genetics 16: e1009162. https://journals.plos.org/plosgenetics/article?id=10.1371/journal.pgen.1009162 (Accessed July 19, 2021).

Buenrostro JD, Giresi PG, Zaba LC, Chang HY, Greenleaf WJ. 2013. Transposition of native chromatin for fast and sensitive epigenomic profiling of open chromatin, DNA-binding proteins and nucleosome position. Nature Methods 2013 10:12 10: 1213–1218. https://www.nature.com/articles/nmeth.2688.

Bürglin TR, Affolter M. 2015. Homeodomain proteins: an update. Chromosoma 2015 125:3 125: 497–521. https://link.springer.com/article/10.1007/s00412-015-0543-8.

Cao J, Spielmann M, Qiu X, Huang X, Ibrahim DM, Hill AJ, Zhang F, Mundlos S, Christiansen L, Steemers FJ, et al. 2019. The single-cell transcriptional landscape of mammalian organogenesis. Nature 2019 566:7745 566: 496–502.

Capellini T, Vaccari G, Ferretti E, Fantini S, He M, Pellegrini M, Quintana L, di Giacomo G, Sharpe J, Selleri L, et al. 2010. Scapula development is governed by genetic interactions of Pbx1 with its family members and with Emx2 via their cooperative control of Alx1. Development 137: 2559–2569. https://pubmed.ncbi.nlm.nih.gov/20627960/.

Capellini TD, di Giacomo G, Salsi V, Brendolan A, Ferretti E, Srivastava D, Zappavigna V, Selleri L. 2006. Pbx1/Pbx2 requirement for distal limb patterning is mediated by the hierarchical control of Hox gene spatial distribution and Shh expression. Development 133: 2263–2273. http://www.ncbi.nlm.nih.gov/pubmed/16672333.

Capellini TD, Handschuh K, Quintana L, Ferretti E, Giacomo G di, Fantini S, Vaccari G, Clarke SL, Wenger AM, Bejerano G, et al. 2011a. Control of pelvic girdle development by genes of the Pbx family and Emx2. Developmental Dynamics 240: 1173–1189. https://pubmed.ncbi.nlm.nih.gov/21455939/.

Capellini TD, Zappavigna V, Selleri L. 2011b. Pbx homeodomain proteins: TALEnted regulators of limb patterning and outgrowth. Developmental Dynamics 240: 1063–1086. http://www.ncbi.nlm.nih.gov/pubmed/21416555.

Capellini TD, Zewdu R, di Giacomo G, Asciutti S, Kugler JE, di Gregorio A, Selleri L. 2008. Pbx1/Pbx2 govern axial skeletal development by controlling Polycomb and Hox in mesoderm and Pax1/Pax9 in sclerotome. Developmental Biology 321: 500–514. http://www.ncbi.nlm.nih.gov/pubmed/18691704.

Chan KKK, Zhang J, Chia NY, Chan YS, Sim HS, Tan KS, Oh SKW, Ng HH, Choo ABH. 2009. KLF4 and PBX1 directly regulate NANOG expression in human embryonic stem cells. Stem cells 27: 2114–2125. https://pubmed.ncbi.nlm.nih.gov/19522013/.

Charite J, McFadden DG, Olson EN. 2000. The bHLH transcription factor dHAND controls Sonic hedgehog expression and establishment of the zone of polarizing activity during limb development. Development 127: 2461–2470.

Chen Y, Gridley T. 2013. Compensatory regulation of the Snai1 and Snai2 genes during chondrogenesis. Journal of bone and mineral research : the official journal of the American Society for Bone and Mineral Research 28: 1412. https://pmc/articles/PMC3663919/.

Chiang C, Litingtung Y, Harris MP, Simandl BK, Li Y, Beachy PA, Fallon JF. 2001. Manifestation of the Limb Prepattern: Limb Development in the Absence of Sonic Hedgehog Function. Developmental Biology 236: 421–435.

Choe SK, Ladam F, Sagerström CG. 2014. TALE factors poise promoters for activation by Hox proteins. Developmental cell 28: 203–211. https://pubmed.ncbi.nlm.nih.gov/24480644/.

Choe SK, Lu P, Nakamura M, Lee J, Sagerström CG. 2009. Meis cofactors control HDAC and CBP accessibility at Hox-regulated promoters during zebrafish embryogenesis. Developmental cell 17: 561–567. https://pubmed.ncbi.nlm.nih.gov/19853569/.

Creuzet S, Couly G, le Douarin NM. 2005. Patterning the neural crest derivatives during development of the vertebrate head: insights from avian studies. Journal of anatomy 207: 447–459. https://pubmed.ncbi.nlm.nih.gov/16313387/.

Darbellay F, Duboule D. 2016. Topological Domains, Metagenes, and the Emergence of Pleiotropic Regulations at Hox Loci. Current topics in developmental biology 116: 299–314.

de Frutos CA, Vega S, Manzanares M, Flores JM, Huertas H, Martínez-Frías ML, Nieto MA. 2007. Snail1 is a transcriptional effector of FGFR3 signaling during chondrogenesis and achondroplasias. Developmental cell 13: 872–883. https://pubmed.ncbi.nlm.nih.gov/18061568/.

de Kumar B, Parker HJ, Paulson A, Parrish ME, Pushel I, Singh NP, Zhang Y, Slaughter BD, Unruh JR, Florens L, et al. 2017. HOXA1 and TALE proteins display cross-regulatory interactions and form a combinatorial binding code on HOXA1 targets. Genome research 27: 1501–1512. https://pubmed.ncbi.nlm.nih.gov/28784834/.

Delgado I, Giovinazzo G, Temiño S, Gauthier Y, Balsalobre A, Drouin J, Torres M. 2021. Control of mouse limb initiation and antero-posterior patterning by Meis transcription factors. Nature Communications 2021 12:1 12: 1–13. https://www.nature.com/articles/s41467-021-23373-9.

Delgado I, López-Delgado AC, Roselló-Díez A, Giovinazzo G, Cadenas V, Fernández-de-Manuel L, Sánchez-Cabo F, Anderson MJ, Lewandoski M, Torres M. 2020. Proximo-distal positional information encoded by an Fgf-regulated gradient of homeodomain transcription factors in the vertebrate limb. Science Advances 6: eaaz0742. https://advances.sciencemag.org/content/6/23/eaaz0742.

DeMare LE, Leng J, Cotney J, Reilly SK, Yin J, Sarro R, Noonan JP. 2013. The genomic landscape of cohesin-associated chromatin interactions. Genome research 23: 1224–1234. http://www.pubmedcentral.nih.gov/articlerender.fcgi?artid=3730097&tool=pmcentrez&rendertype=abstract.

Desanlis I, Paul R, Kmita M. 2020. Transcriptional Trajectories in Mouse Limb Buds Reveal the Transition from Anterior-Posterior to Proximal-Distal Patterning at Early Limb Bud Stage. Journal of Developmental Biology 2020, Vol 8, Page 31 8: 31.

Deschamps J, Duboule D. 2017. Embryonic timing, axial stem cells, chromatin dynamics, and the Hox clock. Genes & development 31: 1406–1416. https://pubmed.ncbi.nlm.nih.gov/28860158/.

di Giacomo G, Koss M, Capellini T, Brendolan A, Pöpperl H, Selleri L. 2006. Spatio-temporal expression of Pbx3 during mouse organogenesis. Gene expression patterns : GEP 6: 747–757. https://pubmed.ncbi.nlm.nih.gov/16434237/.

Dixon JR, Selvaraj S, Yue F, Kim A, Li Y, Shen Y, Hu M, Liu JS, Ren B. 2012. Topological Domains in Mammalian Genomes Identified by Analysis of Chromatin Interactions. Nature 485: 376.

Donaghey J, Thakurela S, Charlton J, Chen JS, Smith ZD, Gu H, Pop R, Clement K, Stamenova EK, Karnik R, et al. 2018. Genetic determinants and epigenetic effects of pioneer-factor occupancy. Nature Genetics 2017 50:2 50: 250–258. https://www.nature.com/articles/s41588-017-0034-3.

Donovan BT, Huynh A, Ball DA, Patel HP, Poirier MG, Larson DR, Ferguson ML, Lenstra TL. 2019. Live-cell imaging reveals the interplay between transcription factors, nucleosomes, and bursting. The EMBO Journal 38: e100809. https://onlinelibrary.wiley.com/doi/full/10.15252/embj.2018100809.

Fabre PJ, Leleu M, Mascrez B, lo Giudice Q, Cobb J, Duboule D. 2018. Heterogeneous combinatorial expression of Hoxd genes in single cells during limb development. BMC Biology 16: 1–15. https://bmcbiol.biomedcentral.com/articles/10.1186/s12915-018-0570-z.

Fernández-Terán MA, Hinchliffe JR, Ros MA. 2006. Birth and death of cells in limb development: A mapping study. Developmental Dynamics 235: 2521–2537.

Ferretti E, Li B, Zewdu R, Wells V, Hebert JM, Karner C, Anderson MJ, Williams T, Dixon J, Dixon MJ, et al. 2011. A Conserved Pbx-Wnt-p63-Irf6 Regulatory Module Controls Face Morphogenesis by Promoting Epithelial Apoptosis. Developmental Cell 21: 627–641.

Galli A, Robay D, Osterwalder M, Bao X, Bénazet J-D, Tariq M, Paro R, Mackem S, Zeller R. 2010. Distinct Roles of Hand2 in Initiating Polarity and Posterior Shh Expression during the Onset of Mouse Limb Bud Development ed. C.J. Tabin. PLoS Genetics 6: e1000901. http://dx.plos.org/10.1371/journal.pgen.1000901.

Garcia-Alonso L, Holland CH, Ibrahim MM, Turei D, Saez-Rodriguez J. 2019. Benchmark and integration of resources for the estimation of human transcription factor activities. Genome research 29: 1363–1375. https://pubmed.ncbi.nlm.nih.gov/31340985/.

Golonzhka O, Nord A, Tang PLF, Lindtner S, Ypsilanti AR, Ferretti E, Visel A, Selleri L, Rubenstein JLR. 2015. Pbx Regulates Patterning of the Cerebral Cortex in Progenitors and Postmitotic Neurons. Neuron 88: 1192–1207.

Grebbin BM, Hau AC, Groß A, Anders-Maurer M, Schramm J, Koss M, Wille C, Mittelbronn M, Selleri L, Schulte D. 2016. PBX1 is required for adult subventricular zone neurogenesis. Development (Cambridge) 143: 2281–2291.

Grebbin BM, Schulte D. 2017. PBX1 as Pioneer Factor: A Case Still Open. Frontiers in Cell and Developmental Biology 5.

Heinz S, Benner C, Spann N, Bertolino E, Lin Y, Laslo P, Cheng J, Murre C, Singh H, Glass C. 2010. Simple combinations of lineage-determining transcription factors prime cis-regulatory elements required for macrophage and B cell identities. Molecular cell 38: 576–589. https://pubmed.ncbi.nlm.nih.gov/20513432/.

Hurtado R, Zewdu R, Mtui J, Liang C, Aho R, Kurylo C, Selleri L, Herzlinger D. 2015. Pbx1-dependent control of VMC differentiation kinetics underlies gross renal vascular patterning. Development (Cambridge) 142: 2653–2664.

Iwafuchi-Doi M, Zaret KS. 2014. Pioneer transcription factors in cell reprogramming. Genes & development 28: 2679–2692. https://pubmed.ncbi.nlm.nih.gov/25512556/.

Jürgens AS, Kolanczyk M, Moebest DCC, Zemojtel T, Lichtenauer U, Duchniewicz M, Gantert MP, Hecht J, Hattenhorst U, Burdach S, et al. 2009. PBX1 is dispensable for neural commitment of RA-treated murine ES cells. In vitro cellular & developmental biology Animal 45: 252–263. https://pubmed.ncbi.nlm.nih.gov/19148706/.

Kelly NH, Huynh NPT, Guilak F. 2020. Single cell RNA-sequencing reveals cellular heterogeneity and trajectories of lineage specification during murine embryonic limb development. Matrix Biology 89: 1–10.

Kmita M, Tarchini B, Zàkàny J, Logan M, Tabin CJ, Duboule D. 2005. Early developmental arrest of mammalian limbs lacking HoxA/HoxD gene function. Nature 2005 435:7045 435: 1113–1116. https://www.nature.com/articles/nature03648.

Koss M, Bolze A, Brendolan A, Saggese M, Capellini TD, Bojilova E, Boisson B, Prall OWJ, Elliott DA, Solloway M, et al. 2012. Congenital Asplenia in Mice and Humans with Mutations in a Pbx/Nkx2-5/p15 Module. Developmental Cell 22: 913–926.

Kothary R, Clapoff S, Darling S, Perry MD, Moran LA, Rossant J. 1989. Inducible expression of an hsp68-lacZ hybrid gene in transgenic mice. Development 105: 707–714. https://pubmed.ncbi.nlm.nih.gov/2557196/.

Kozhemyakina E, Ionescu A, Lassar AB. 2014. GATA6 Is a Crucial Regulator of Shh in the Limb Bud. PLOS Genetics 10: e1004072. https://journals.plos.org/plosgenetics/article?id=10.1371/journal.pgen.1004072.

Kuijper S, Beverdam A, Kroon C, Brouwer A, Candille S, Barsh G, Meijlink F. 2005. Genetics of shoulder girdle formation: roles of Tbx15 and aristaless-like genes. Development 132: 1601–1610.

Kvon EZ, Zhu Y, Kelman G, Visel A, Dickel DE, Pennacchio LA. 2020. Comprehensive In Vivo Interrogation Reveals Phenotypic Impact of Human Enhancer Variants. Cell 180. https://doi.org/10.1016/j.cell.2020.02.031.

Lee TI, Young RA. 2013. Transcriptional Regulation and its Misregulation in Disease. Cell 152: 1237. https://pmc/articles/PMC3640494/.

Lettice LA, Williamson I, Devenney PS, Kilanowski F, Dorin J, Hill RE. 2014. Development of five digits is controlled by a bipartite long-range cis-regulator. Development 141: 1715–1725.

Lettice LA, Williamson I, Wiltshire JH, Peluso S, Devenney PS, Hill AE, Essafi A, Hagman J, Mort R, Grimes G, et al. 2012. Opposing Functions of the ETS Factor Family Define Shh Spatial Expression in Limb Buds and Underlie Polydactyly. Developmental Cell 22: 459. https://pmc/articles/PMC3314984/.

Lickwar CR, Mueller F, Hanlon SE, McNally JG, Lieb JD. 2012. Genome-wide protein–DNA binding dynamics suggest a molecular clutch for transcription factor function. Nature 2012 484:7393 484: 251–255. https://www.nature.com/articles/nature10985.

Liu Y, Ma L, Wu L, Luo W, Kundu R, Sangiorgi F, Snead M, Maxson R. 1994. Regulation of the Msx2 homeobox gene during mouse embryogenesis: a transgene with 439 bp of 5’ flanking sequence is expressed exclusively in the apical ectodermal ridge of the developing limb. Mechanisms of development 48: 187–197. https://pubmed.ncbi.nlm.nih.gov/7893602/.

Losa M, Latorre V, Andrabi M, Ladam F, Sagerström C, Novoa A, Zarrineh P, Bridoux L, Hanley NA, Mallo M, et al. 2017. A tissue-specific, Gata6-driven transcriptional program instructs remodeling of the mature arterial tree. eLife 6.

Losa M, Risolino M, Li B, Hart J, Quintana L, Grishina I, Yang H, Choi IF, Lewicki P, Khan S, et al. 2018. Face morphogenesis is promoted by Pbx-dependent EMT via regulation of Snail1 during frontonasal prominence fusion. Development 145.

Lowe LA, Yamada S, Kuehn MR. 2000. HoxB6-Cre Transgenic Mice Express Cre Recombinase in Extra-Embryonic Mesoderm, in Lateral Plate and Limb Mesoderm and at the Midbrain/Hindbrain Junction. Genesis 118–120.

Malkmus J, Ramos Martins L, Jhanwar S, Kircher B, Palacio V, Sheth R, Leal F, Duchesne A, Lopez-Rios J, Peterson KA, et al. 2021. Spatial regulation by multiple Gremlin1 enhancers provides digit development with cis-regulatory robustness and evolutionary plasticity. Nature communications 12. https://pubmed.ncbi.nlm.nih.gov/34548488/.

Mann RS, Lelli KM, Joshi R. 2009. Hox specificity unique roles for cofactors and collaborators. Current topics in developmental biology 88: 63–101. https://pubmed.ncbi.nlm.nih.gov/19651302/.

Mao J, McGlinn E, Huang P, Tabin CJ, McMahon AP. 2009. Fgf-Dependent Etv4/5 Activity Is Required for Posterior Restriction of Sonic hedgehog and Promoting Outgrowth of the Vertebrate Limb. Developmental Cell 16: 600–606.

Mariani L, Guo X, Menezes NA, Drozd AM, Çakal SD, Wang Q, Ferretti E. 2021. A TALE/HOX code unlocks WNT signalling response towards paraxial mesoderm. Nature communications 12. https://pubmed.ncbi.nlm.nih.gov/34446717/.

McCulley DJ, Wienhold MD, Hines EA, Hacker TA, Rogers A, Pewowaruk RJ, Zewdu R, Chesler NC, Selleri L, Sun X. 2018. PBX transcription factors drive pulmonary vascular adaptation to birth. Journal of Clinical Investigation 128: 655–667.

McLean C, Bristor D, Hiller M, Clarke S, Schaar B, Lowe C, Wenger A, Bejerano G. 2010. GREAT improves functional interpretation of cis-regulatory regions. Nature biotechnology 28: 495–501. https://pubmed.ncbi.nlm.nih.gov/20436461/.

Mirny LA. 2010. Nucleosome-mediated cooperativity between transcription factors. Proceedings of the National Academy of Sciences of the United States of America 107: 22534–22539. https://www.pnas.org/content/107/52/22534.

Moens CB, Selleri L. 2006. Hox cofactors in vertebrate development. Developmental Biology.

Monti R, Barozzi I, Osterwalder M, Lee E, Kato M, Garvin TH, Plajzer-Frick I, Pickle CS, Akiyama JA, Afzal V, et al. 2017. Limb-Enhancer Genie: An accessible resource of accurate enhancer predictions in the developing limb. PLoS Computational Biology 13: e1005720. https://pmc/articles/PMC5578682/.

Morgunova E, Taipale J. 2017. Structural perspective of cooperative transcription factor binding. Current Opinion in Structural Biology 47: 1–8.

Moyle-Heyrman G, Tims HS, Widom J. 2011. Structural constraints in collaborative competition of transcription factors against the nucleosome. Journal of molecular biology 412: 634–646. https://pubmed.ncbi.nlm.nih.gov/21821044/.

Nakamura E, Hata K, Takahata Y, Kurosaka H, Abe M, Abe T, Kihara M, Komori T, Kobayashi S, Murakami T, et al. 2021. Zfhx4 regulates endochondral ossification as the transcriptional platform of Osterix in mice. Communications Biology 2021 4:1 4: 1–11. https://www.nature.com/articles/s42003-021-02793-9.

Nguyen M, Zhu J, Nakamura E, Bao X, Mackem S. 2009. Tamoxifen-dependent, inducible Hoxb6CreERT recombinase function in lateral plate and limb mesoderm, CNS isthmic organizer, posterior trunk neural crest, hindgut, and tailbud. Developmental dynamics : an official publication of the American Association of Anatomists 238: 467–474. https://pubmed.ncbi.nlm.nih.gov/19161221/.

Osterwalder M, Speziale D, Shoukry M, Mohan R, Ivanek R, Kohler M, Beisel C, Wen X, Scales SJ, Christoffels VM, et al. 2014. HAND2 targets define a network of transcriptional regulators that compartmentalize the early limb bud mesenchyme. Developmental cell 31: 345–357. http://www.ncbi.nlm.nih.gov/pubmed/25453830.

Osterwalder M, Tran S, Hunter RD, Meky EM, von Maydell K, Harrington AN, Godoy J, Novak CS, Plajzer-Frick I, Zhu Y, et al. 2022. Characterization of Mammalian In Vivo Enhancers Using Mouse Transgenesis and CRISPR Genome Editing. In Methods in Molecular Biology, Vol. 2403 of, pp. 147–186, Humana, New York, NY https://link.springer.com/protocol/10.1007/978-1-0716-1847-9_11.

Parker HJ, de Kumar B, Green SA, Prummel KD, Hess C, Kaufman CK, Mosimann C, Wiedemann LM, Bronner ME, Krumlauf R. 2019. A Hox-TALE regulatory circuit for neural crest patterning is conserved across vertebrates. Nature communications 10. https://pubmed.ncbi.nlm.nih.gov/30867425/.

Penkov D, SanMartín DM, Fernandez-Díaz LC, Rosselló CA, Torroja C, Sánchez-Cabo F, Warnatz HJ, Sultan M, Yaspo ML, Gabrieli A, et al. 2013. Analysis of the DNA-Binding Profile and Function of TALE Homeoproteins Reveals Their Specialization and Specific Interactions with Hox Genes/Proteins. Cell Reports 3: 1321–1333.

Pöpperl H, Bienz M, Studer M, Chan SK, Aparicio S, Brenner S, Mann RS, Krumlauf R. 1995. Segmental expression of Hoxb-1 is controlled by a highly conserved autoregulatory loop dependent upon exd/pbx. Cell 81: 1031–1042. https://pubmed.ncbi.nlm.nih.gov/7600572/.

Rhee JW, Arata A, Selleri L, Jacobs Y, Arata S, Onimaru H, Cleary ML. 2004. Pbx3 Deficiency Results in Central Hypoventilation. The American Journal of Pathology 165: 1343. https://pmc/articles/PMC1618620/.

Royle S, Tabin C, Young J. 2021. Limb positioning and initiation: An evolutionary context of pattern and formation. Developmental dynamics : an official publication of the American Association of Anatomists.

Ryoo HD, Mann RS. 1999. The control of trunk Hox specificity and activity by Extradenticle. Genes & development 13: 1704–1716. https://pubmed.ncbi.nlm.nih.gov/10398683/.

Sagai T, Hosoya M, Mizushina Y, Tamura M, Shiroishi T. 2005. Elimination of a long-range cis-regulatory module causes complete loss of limb-specific Shh expression and truncation of the mouse limb. Development 132: 797–803. http://www-gsd.lbl.gov/vista/index.shtml.

Sagerström CG. 2004. PbX marks the spot. Developmental cell 6: 737–738. https://pubmed.ncbi.nlm.nih.gov/15177017/.

Selleri L, Depew MJ, Jacobs Y, Chanda K, Tsang KY, Cheah KSE, Rubenstein JLR, O’Gorman S, Cleary ML. 2001. Requirement for Pbx1 in skeletal patterning and programming chondrocyte proliferation and differentiation. Development 128: 3543–3557.

Selleri L, DiMartino J, van Deursen J, Brendolan A, Sanyal M, Boon E, Capellini T, Smith KS, Rhee J, Pöpperl H, et al. 2004. The TALE Homeodomain Protein Pbx2 Is Not Essential for Development and Long-Term Survival. Molecular and Cellular Biology 24: 5324–5331. https://pubmed.ncbi.nlm.nih.gov/15169896/.

Selleri L, Zappavigna V, Ferretti E. 2019. ‘Building a perfect body’: Control of vertebrate organogenesis by PBX-dependent regulatory networks. Genes and Development 33: 258–275. https://pubmed.ncbi.nlm.nih.gov/30824532/.

Sonnet W, Rezsöhazy R, Donnay I. 2012. Characterization of TALE genes expression during the first lineage segregation in mammalian embryos. Developmental dynamics : an official publication of the American Association of Anatomists 241: 1827–1839. https://pubmed.ncbi.nlm.nih.gov/22987645/.

Stankunas K, Shang C, Twu KY, Kao S-C, a Jenkins N, Copeland NG, Sanyal M, Selleri L, Cleary ML, Chang C-P. 2008. Pbx/Meis deficiencies demonstrate multigenetic origins of congenital heart disease. Circulation research 103: 702–709.

Sun X, Lewandoski M, Meyers EN, Liu Y-H, Maxson RE, Martin GR. 2000. Conditional inactivation of Fgf4 reveals complexity of signalling during limb bud development. Nature Genetics 2000 25:1 25: 83–86. https://www-nature-com.ucsf.idm.oclc.org/articles/ng0500_83.

te Welscher P, Fernandez-Teran M, Ros MA, Zeller R. 2002. Mutual genetic antagonism involving GLI3 and dHAND prepatterns the vertebrate limb bud mesenchyme prior to SHH signaling. Genes & Development 16: 421–426. http://genesdev.cshlp.org/content/16/4/421.full.

Tzchori I, Day TF, Carolan PJ, Zhao Y, Wassif CA, Li LQ, Lewandoski M, Gorivodsky M, Love PE, Porter FD, et al. 2009. LIM homeobox transcription factors integrate signaling events that control three-dimensional limb patterning and growth. Development 136: 1375–1385.

Wagner K, Mincheva A, Korn B, Lichter P, Pöpperl H. 2001. Pbx4, a new Pbx family member on mouse chromosome 8, is expressed during spermatogenesis. Mechanisms of Development 103: 127–131.

Welsh IC, Hart J, Brown JM, Hansen K, Marques MR, Aho RJ, Grishina I, Hurtado R, Herzlinger D, Ferretti E, et al. 2018. Pbx loss in cranial neural crest, unlike in epithelium, results in cleft palate only and a broader midface. Journal of Anatomy.

Xu B, Wellik DM. 2011. Axial Hox9 activity establishes the posterior field in the developing forelimb. Proceedings of the National Academy of Sciences of the United States of America 108: 4888–4891. https://pubmed.ncbi.nlm.nih.gov/21383175/.

Yin W, Mendoza L, Monzon-Sandoval J, Urrutia AO, Gutierrez H. 2021. Emergence of co-expression in gene regulatory networks. PLOS ONE 16: e0247671. https://journals.plos.org/plosone/article?id=10.1371/journal.pone.0247671.

Zuniga A, Zeller R. 2020. Dynamic and self-regulatory interactions among gene regulatory networks control vertebrate limb bud morphogenesis. Current topics in developmental biology 139: 61–88. https://pubmed.ncbi.nlm.nih.gov/32450969/.

