## Supplementary material for "A spatio-temporally constrained gene regulatory network directed by PBX1/2 acquires limb patterning specificity via HAND2": Supplmentary files

#### SUPPLEMENTARY MATERIALS

#### SUPPL. FIGURE LEGENDS

**Supplementary Figure 1. Absence of limb and overall skeletal abnormalities in embryos with *Pbx1* conditional deletion in MSX2-cells on a *Pbx2*-deficient background.**

(A-C) Time-course by IF showing PBX1 protein (red) localization (red arrows) in developing hindlimbs from E11.0 to E12.0. DAPI labels nuclei (blue). (D-G) Cartilage preparations from E14.5 whole embryos and hindlimb autopods with conditional deletion of *Pbx1* on a *Pbx2*-deficient background using a *Msx2Cre* driver, compared with their respective littermate controls, as assessed by Alcian Blue staining. (H-I') Deletion of PBX1 protein in developing hindlimb buds assessed by immunohistochemistry with a specific PBX1 antibodies. Consistent with activity of *Msx2Cre* (Liu et al. 1994), PBX1 is lost from the ventral hindlimb apical ectodermal ridge (AER) and ventral ectoderm of E10.25 $Pbx1^{f/f};Pbx2^{+/-};Msx2^{Cre/+}$ mutant embryos (I,I') compared to controls (H,H'). (J) Ratio of Tibia/Femur length in mutant $Pbx1^{f/f};Pbx2^{-/-};Hoxb6^{CreERT/+}$ embryos (Mut) compared to control $Pbx1^{f/f};Pbx2^{-/-}$ embryos (Co). E9.5, E10.0 and E10.5 indicate the timepoints of tamoxifen injections.

**Supplementary Figure 2. Annotation of fine-grained cell clusters in E10.5 hindlimb bud based on top differentially expressed markers.**

(A) UMAP representation of 9,859 hindlimb cells in scRNAseq of E10.5 hindlimb buds annotated with clusters at fine-grained resolution. (B) Dot plot of the highest differentially expressed marker genes for each of the cluster shown in (A). The size of each dot represents the proportion of cells within a given population that expresses the gene; the intensity of color indicates the average level of expression of the indicated gene. (C) UMAP (color-coded based on the expression level of the genes; right) and violin plot (left) of normalized expression of *Pbx2*.

**Supplementary Figure 3. Top differentially expressed genes for each cell cluster identified by scRNAseq in E10.5 hindlimb bud.**

(A-F) UMAPs (color-coded based on the expression level of the genes) of the most differentially expressed marker genes identified for each coarse-grained cell cluster (see Fig. 2B): Erythrocytes (*Hba-a2, Klf1*); Endothelial cells (*Pecam1, Emcn*); Epithelial cells (*Wnt6, Epcam*); Phagocytes (*Spi1, Fcer1g*); and pluripotent progenitors (*Sox2, Oct4*). (G) UMAP color-coded based on the expression level of mesenchymal-specific markers as well as *Hox* genes, in mesenchymal hindlimb bud cells at E10.5.

**Supplementary Figure 4. Genome-wide analysis of PBX1, HAND2, and PBX1-HAND2 binding and chromatin landscapes in E10.5 mouse hindlimb buds.**

(A, B) Bar plots showing the distribution of peaks bound by PBX1-only, HAND2-only, or co-bound (Both) relative to genomic regions (Intergenic, Promoter, or Intragenic regions). (C) Table summarizing the number of replicated peaks and the final coverage for all generated datasets. All datasets were generated in this study, except for the published CTCF ChIPseq (DeMare et al. 2013). (D-G) Top over-represented biological processes (D,F) and mouse phenotypes (E,G) associated to PBX1-only and HAND2-only peaks, respectively. Values on the x axis correspond to the binomial (uncorrected) *p*-value of the enrichment. (H) Heatmap highlighting the enrichment of known transcription factor motifs in the regions bound by PBX1-HAND2 (Both), PBX1-only, or HAND2-only. Five subgroups are highlighted. The top bracket corresponds to transcription factor motifs correlated to PBX1 binding, mainly at TSS-distal domains and associated to homeobox transcription factors. The second bracket corresponds to transcription factor motifs enriched at TSS-distal sites and comprising bHLH and homeobox transcription factors. The third bracket includes PBX1-specific cofactors, such as PREP1 and PBX3. The last bracket comprises SMAD transcription factor motifs, which are enriched at TSS-distal regions.

**Supplementary Figure 5. Correlation of chromatin state with the peaks bound by PBX1-only or HAND2-only.**

(A) Distribution of intergenic regions bound by HAND2 alone, relative to their distance to the nearest H3K27ac peak, H3K27me3-enriched region, or accessible region (as measured by ATACseq). (B) Same as (A), including intergenic regions bound by PBX1 alone. (C-D) Same as (A), including regions bound by either HAND2 or PBX1 alone, at promoters.

**Supplementary Figure 6. Comprehensive analysis of PBX1-bound *Hand2* limb enhancers via *LacZ* transgenic reporter assays.**

(A-Q) Representative mouse embryos from *LacZ* transgenic reporter assays for assessment of *in vivo* enhancer activities at E10.5 and/or E11.5. Enhancer activity domains appear blue due to X-gal staining. Vista enhancer IDs (mm numbers) are listed on top of each panel and the relative distance of the enhancer to the *Hand2* TSS is indicated (in kb). Numbers at the bottom right of each embryo indicate the tissue-specific reproducibility of the *LacZ* signal(s). NRS: no reproducible staining.

**Supplementary Figure 7. Analyses of *Pbx1/2* and *Hand2* mutant hindlimb buds transcriptomes.**

(A) Heatmap showing the relative expression (z-score) of DEGs identified by comparing the transcriptomes of wild-type (Ctrl) and *Pbx1/2* mutant (top left) or *Hand2* mutant (top right) mouse embryonic hindlimb buds at E10.5. DEGs are genes whose expression is significantly changed between Ctrl and mutant samples (Fold Change equal or larger than 2, in either direction; FDR <= 0.05). Box plots showing the counts per million (CPM) of *Pbx1* (bottom left) and *Hand2* (bottom right) transcripts from the respective bulk RNASeq experiments. Biological replicates in each box plot indicated by red dots. (B) Table showing the numbers of upregulated (Up) and downregulated (Down) DEGs in *Pbx1/2* (*Pbx*) mutant and *Hand2* mutant hindlimb buds, based on the genes that were robustly detected in both sets of experiments. (C) Summary of Biological Processes GO terms associated with genes up-regulated, down-regulated and discordant in hindlimb buds of *Pbx1/2* mutant, *Hand2* mutant, or both, compared to controls. GO term categories are separated also based on binding by PBX1 only, HAND2 only or Both as determined by ChIPseq. (D) UMAP representation of 9,859 hindlimb cells in scRNAseq of E10.5 hindlimb buds color-coded according to the enrichment of *Pbx1/2* and target genes using an available compendium (Dorothea, Garcia-Alonso et al. 2019)). Analysis shows that *Pbx1/2* targets exhibit highest enrichment in mesenchymal subclusters 7 and 12 (highlighted in orange), which correspond to the two clusters showing also the highest enrichment of *Hand2* target genes.

**Supplementary Figure 8. Integration of chromatin and transcriptomic datasets.**
Each TAD was scored based on the number of total sites bound by PBX1 and/or HAND2 within it, and the strength of the individual binding site (ChIPseq enrichment). For each TAD, the scores were integrated separately for three groups (bound by PBX1-only, HAND2-only, or both). For inter-TAD comparison, the scores were re-scaled based on the number of expressed genes in the TAD. Each gene was assigned the scores of the TAD and the genes were separated based on RNAseq classification (X axis of the plots).

**Supplementary Table 1. List of the total number of embryos analyzed with *Pbx1* conditional deletion in MSX2- and HOXB6-positive cells on a *Pbx2*-deficient background.**

**Supplementary Table 2. Complete lists of replicated ChIPseq peaks identified in wild-type mouse limbs for PBX1, HAND2, H3K27ac and H3K27me3, as well as of replicated accessible regions as defined by ATACseq**. Coordinates (chromosome, start and end) and combined *p*-value (as -log10) are shown.

**Supplementary Table 3. List of *Hand2* enhancer elements tested via LacZ transgenic reporter assays.** Coordinates and distance to the *Hand2* TSS are indicated (mm10).

**Supplementary Table 4. Lists of DEGs in *Pbx1cKO$^{Mes}$;Pbx2$^{-/-}$* and *Hand2cKO$^{Mes}$* hindlimb buds compared to their respective littermate controls and intersection of RNAseq and ChIPseq datasets.** *Pbx_mutant* lists the results of the RNAseq data analysis performed with edgeR comparing *Pbx1cKO$^{Mes}$;Pbx2$^{-/-}$* mutant to control hindlimb buds. The table shows, for each gene, the counts per million (CPM) for each replicate, along with the log2-fold-change, the *p*-value and the FDR. The last two columns indicate whether the gene was defined as up- or down-regulated in *Pbx* mutant hindlimb buds. *Hand_mutant* lists the results of the RNAseq analysis on *Hand2cKO$^{Mes}$* mutant hindlimb buds. The content of this table is as described in *Pbx_mutant*. *Summary* lists the RNAseq results, in which each gene has also been annotated as direct PBX1 and/or HAND2 target based on intersection with the results of the ChIPseq. For each gene, the table shows the log2-fold-change and FDR, separately for the *Pbx* and *Hand2* mutant hindlimb transcriptomes, along with columns indicating whether the gene was defined as significantly up- or down-regulated. *Regulation* lists the classification of each gene based on the behavior in the RNAseq profiles. The last six columns indicate whether the gene is predicted as a direct PBX1 or HAND2 target, based on the peaks identified via ChIPseq, either in the proximity of the gene promoter (within 2.5 kbp) or within 10 or 100 Kbp from the main annotated TSS of the gene. *Summary_Numbers* illustrates the breakdown of the DEG counts in the mutant limbs, based on how they have been classified (see *Summary*). Summary_100kbp and Summary_10kbp show the list of the total number of ChIPseq peaks near target genes, defined by our RNAseq datasets, within a 100kb and 10kb window, respectively.

**Supplementary Table 5. List of numbers of replicated peaks for each ChIPseq dataset from hindlimb (HL), midface (MF) and branchial arch 2 (BA2), respectively, and numbers of overlapping peaks for each intersection.**

#### SUPPL. MATERIALS AND METHODS

##### Enhancer Mutagenesis in the Mouse

For coordinates of enhancer mm1828 and mm1689, see Supplementary Table 2. The PBX and PBX-HOX binding motifs that were mutagenized follow below:

<u>Enhancer mm1689</u> contained 7 PBX or PBX-HOX binding sites at the base-pair positions 703, 1059, 1366, 1482, 1560, 1765 and 1902.

In all cases we disrupted the binding sites by substituting T/A bases into C and C/G bases into A.

##### scRNAseq data analysis

The Cell Ranger v2.2.0 pipeline from 10X Genomics was used for initial processing of raw sequencing reads. Briefly, raw sequencing reads were demultiplexed, aligned to the mouse genome (mm10), filtered for quality using default parameters, and UMI counts were calculated for each gene per cell. Filtered gene-barcode matrices were then analyzed using the Seurat v3.0 R package(Stuart et al. 2019). Cells were filtered to ensure that only those showing a number of total expressed transcripts between 3,000 and 25,000, corresponding to at least 1,000 expressed genes, and with the mass of transcripts derived from the mitochondrial chromosomes representing less than 10% of the total. Data were normalized using scTransform (Hafemeister and Satija 2019), using the best 5,000 features. Cell clusters were identified by constructing a shared nearest neighbor graph followed by a modularity optimization-based clustering algorithm (Leiden algorithm) using the top 60 principal components as determined by PC. Clustering was performed at multiple resolutions between 0.2 and 2, and optimal resolution was determined empirically based on the expression of known population markers (resolution = 0.8). Cells were visualized in two-dimensional space using Uniform Manifold Approximation and Projection (UMAP) dimensional reduction. Markers for each cluster were identified using the FindAllMarkers function using the Wilcoxon test and setting the *min.pct* to 0.1 and *logfc.threshold* to 0.25. Cluster identity was manually annotated based on the expression of known marker genes. To identify how well each target gene was co-expressed with either *Pbx1/2*, *Hand2*, or both, we focused on the mesenchymal clusters and used as a statistical threshold an 'area under the curve' (AUC) >=0.55. Dorothea (Garcia-Alonso et al. 2019) was used to retrieve the computationally predicted target genes of PBX1/2 and HAND2. Then these lists of genes were used to score each single-cell for their overall expression, using the function AddModuleScore from Seurat v3.0.

##### Bulk RNAseq data analysis

Reads for each tissue were mapped against the mouse genome (mm10) using the Tophat 2 aligner (version 2.0.13) with default parameter settings except for setting the flag --no-coverage-search. Expression levels for each tissue were initially quantified using htseq-count. Differential expression analyses (*Pbx1*$^{f/f}$*;Pbx2*$^{-/-}$*;HoxB6*$^{Cre/+}$ vs littermate controls; and *Hand2*$^{f/f}$*;HoxB6*$^{Cre/+}$ vs littermate controls) was performed using the Bioconductor package edgeR (Robinson et al. 2010). Briefly, after estimating global and gene-wise dispersion parameters, normalization was performed using TMM (trimmed mean of M-vaues) (Robinson and Oshlack 2010) Genes with a fold change >= 1.2 or <= −1.2 and a FDR <= 0.05 were defined as differentially expressed genes (DEGs).

**ChIPseq data analysis**

ChIPseq reads were aligned to the mm10 release of the mouse genome (Dec. 2011, GRCm38) using Bowtie (Langmead and Salzberg 2012) with parameters -v 2 -m 1. Peak calling was performed using Model-Based Analysis for ChIPseq (MACS) v1.4 (Zhang et al., 2008) with matched input DNA as control and parameters --gsize=mm --bw=150 --nomodel --shiftsize=100. Each experiment was performed in duplicate; for HAND2 and PBX1 ChIPseq analysis, peaks detected in both replicates were merged using a statistical method that takes into account the combined statistical evidence from the two replicates (MSPC; parameters: -r biological -s 1E-10 -W 1E-6, Jalili et al. 2015). Briefly, ChIPseq peaks were associated to genes using custom scripts. Hypergeometric Optimization of Motif EnRichment (HOMER, Heinz et al. 2010) was used to perform enrichment analysis for known transcription-factor-binding sites as well as de novo motif discovery.

#### SUPPLEMENTARY REFERENCES


DeMare LE, Leng J, Cotney J, Reilly SK, Yin J, Sarro R, Noonan JP. 2013. The genomic landscape of cohesin-associated chromatin interactions. *Genome research* **23**: 1224–1234. http://www.pubmedcentral.nih.gov/articlerender.fcgi?artid=3730097&tool=pmcentrez&rendertype=abstract.

Garcia-Alonso L, Holland CH, Ibrahim MM, Turei D, Saez-Rodriguez J. 2019. Benchmark and integration of resources for the estimation of human transcription factor activities. *Genome research* **29**: 1363–1375. https://pubmed.ncbi.nlm.nih.gov/31340985/

Hafemeister C, Satija R. 2019. Normalization and variance stabilization of single-cell RNA-seq data using regularized negative binomial regression. *Genome Biology 2019 20:1* **20**: 1–15. https://genomebiology.biomedcentral.com/articles/10.1186/s13059-019-1874-1.

Heinz S, Benner C, Spann N, Bertolino E, Lin Y, Laslo P, Cheng J, Murre C, Singh H, Glass C. 2010. Simple combinations of lineage-determining transcription factors prime cis-regulatory elements required for macrophage and B cell identities. *Molecular cell* **38**: 576–589. https://pubmed.ncbi.nlm.nih.gov/20513432/

Jalili V, Matteucci M, Masseroli M, Morelli MJ. 2015. Using combined evidence from replicates to evaluate ChIP-seq peaks. *Bioinformatics (Oxford, England)* **31**: 2761–2769. https://pubmed.ncbi.nlm.nih.gov/25957351/

Langmead B, Salzberg SL. 2012. Fast gapped-read alignment with Bowtie 2. *Nature Methods 2012 9:4* **9**: 357–359. https://www.nature.com/articles/nmeth.1923

Liu Y, Ma L, Wu L, Luo W, Kundu R, Sangiorgi F, Snead M, Maxson R. 1994. Regulation of the Msx2 homeobox gene during mouse embryogenesis: a transgene with 439 bp of 5' flanking sequence is expressed exclusively in the apical ectodermal ridge of the developing limb. *Mechanisms of development* **48**: 187–197. https://pubmed.ncbi.nlm.nih.gov/7893602/

Robinson MD, McCarthy DJ, Smyth GK. 2010. edgeR: a Bioconductor package for differential expression analysis of digital gene expression data. *Bioinformatics (Oxford, England)* **26**: 139–140. https://pubmed.ncbi.nlm.nih.gov/19910308/

Robinson MD, Oshlack A. 2010. A scaling normalization method for differential expression analysis of RNA-seq data. *Genome biology* **11**. https://pubmed.ncbi.nlm.nih.gov/20196867/

Stuart T, Butler A, Hoffman P, Hafemeister C, Papalexi E, Mauck WM, Hao Y, Stoeckius M, Smibert P, Satija R. 2019. Comprehensive Integration of Single-Cell Data. *Cell* **177**: 1888-1902.e21. http://www.cell.com/article/S0092867419305598/fulltext

### Supplementary Figure 1

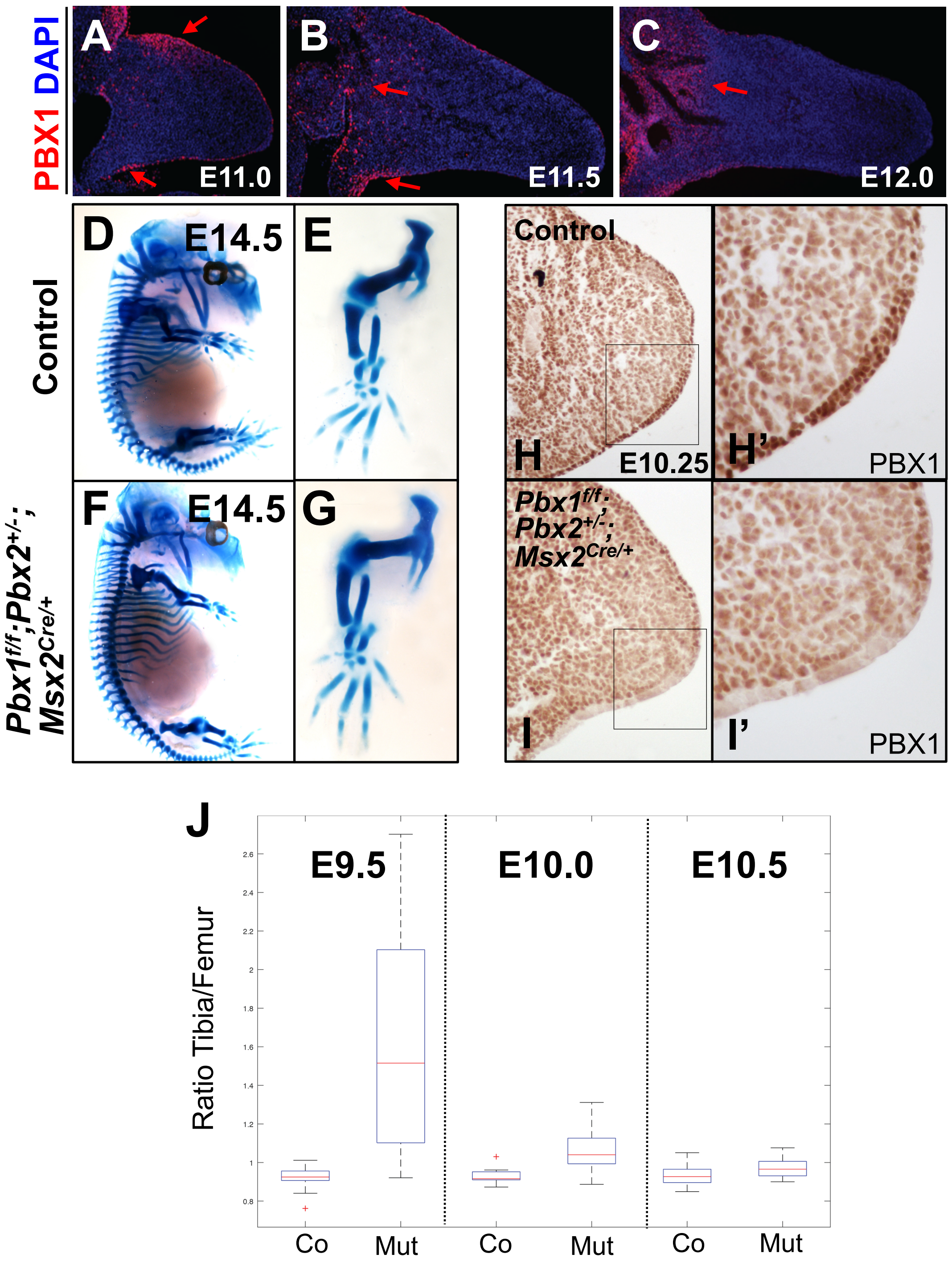

### Supplementary Figure 2

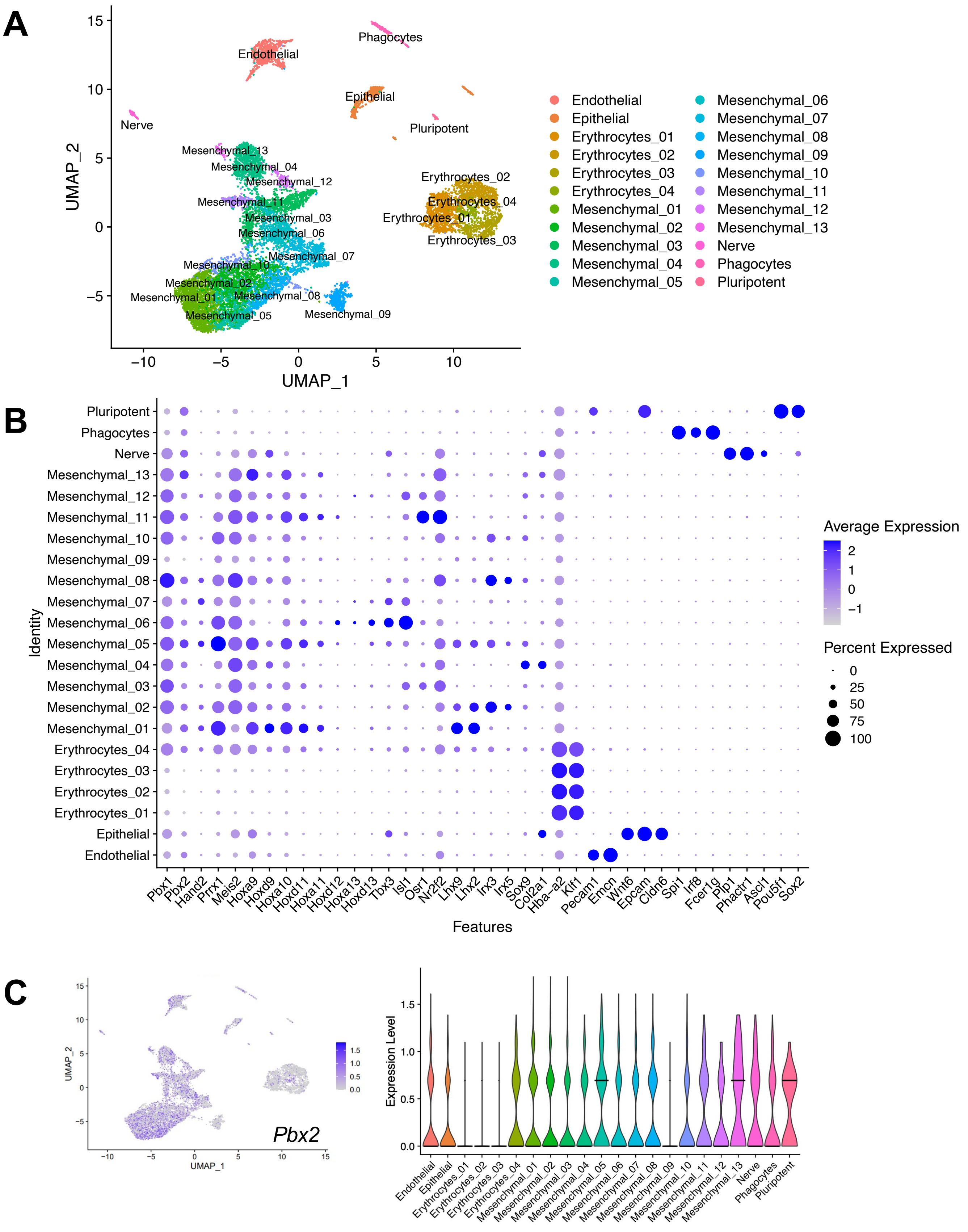

### Supplementary Figure 3

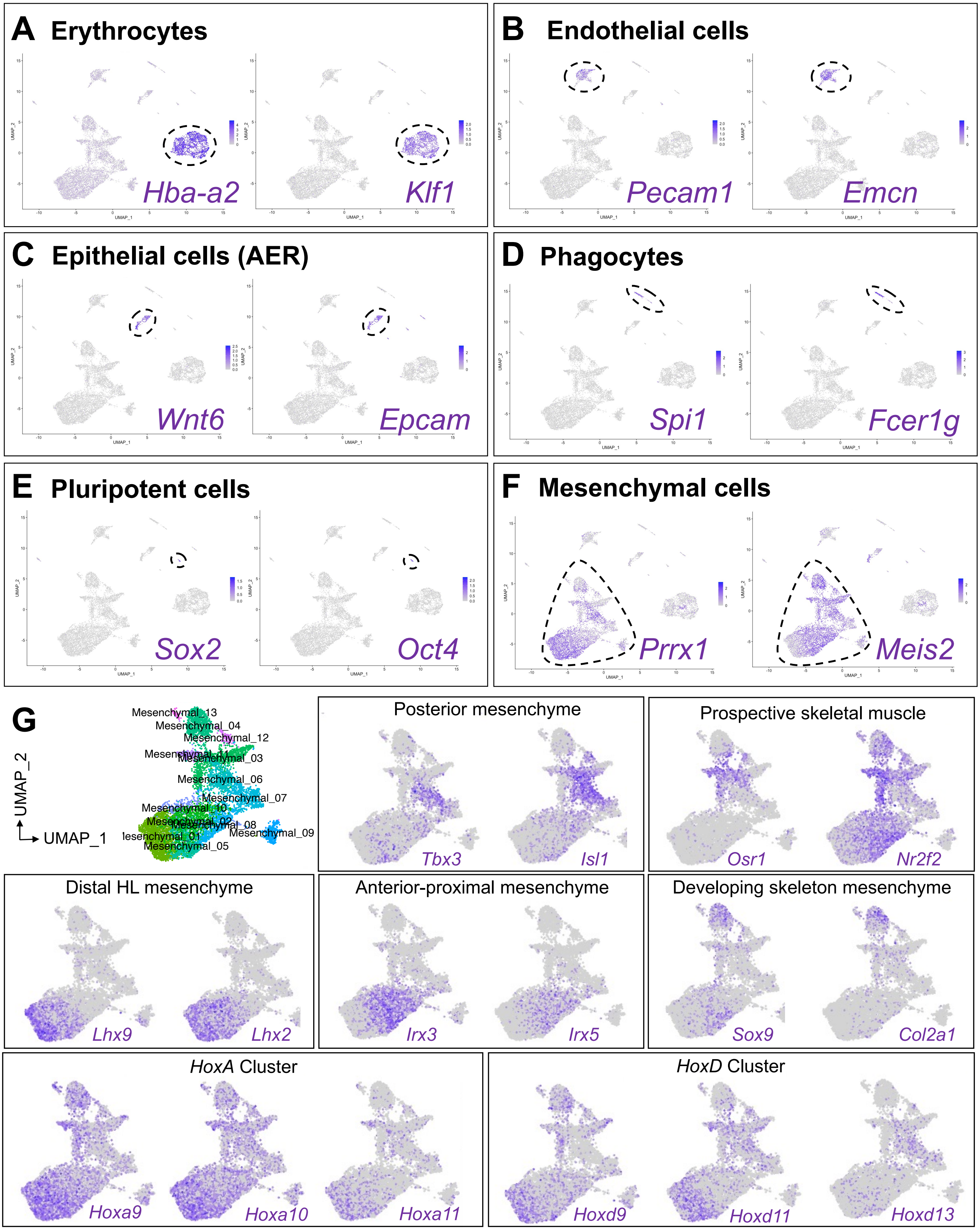

### Supplementary Figure 4

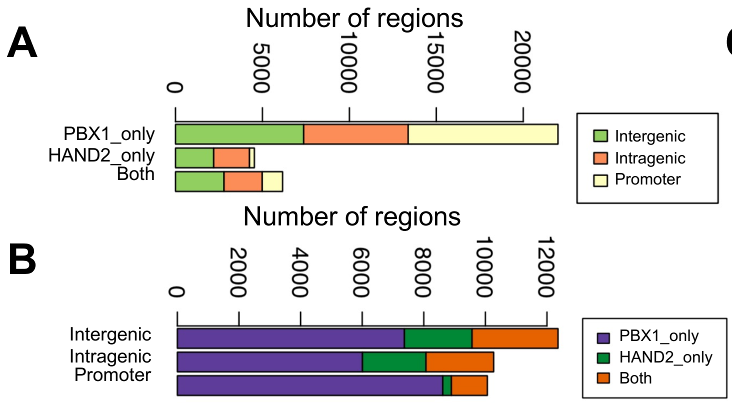


C

| | Replicated peaks | Coverage (bp) |
|---|---|---|
| ATAC-seq | 24,795 | 34,030,209 |
| PBX1 ChIP-seq | 28,177 | 43,497,077 |
| HAND2 ChIP-seq | 10,820 | 11,182,158 |
| H3K27ac ChIP-seq | 33,898 | 307,330,502 |
| H3K27me3 ChIP-seq | 9,866 | 89,606,534 |
| CTCF ChIP-seq (*) | 15,046 | 15,820,050 |

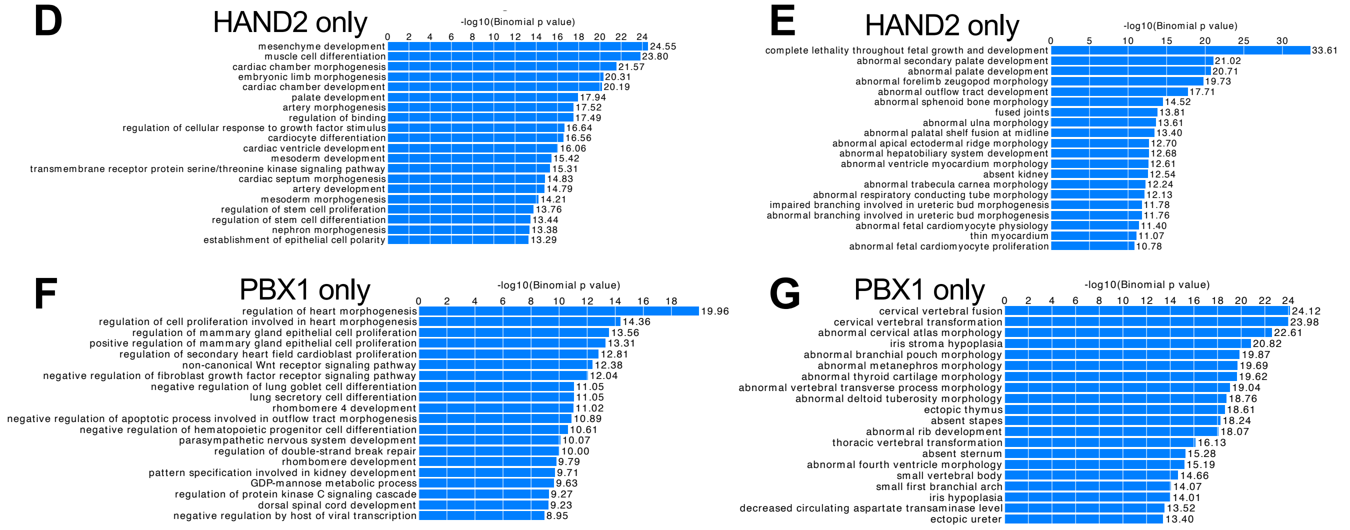


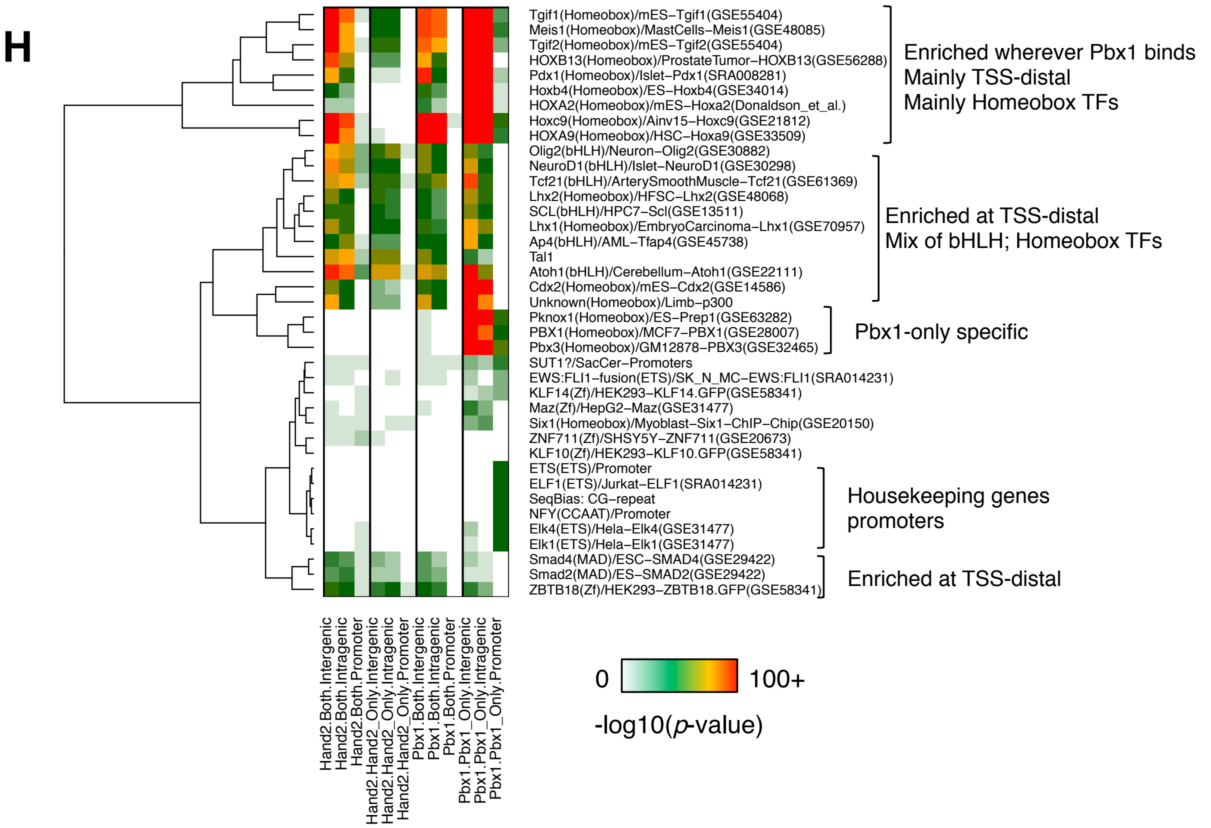

### Supplementary Figure 5

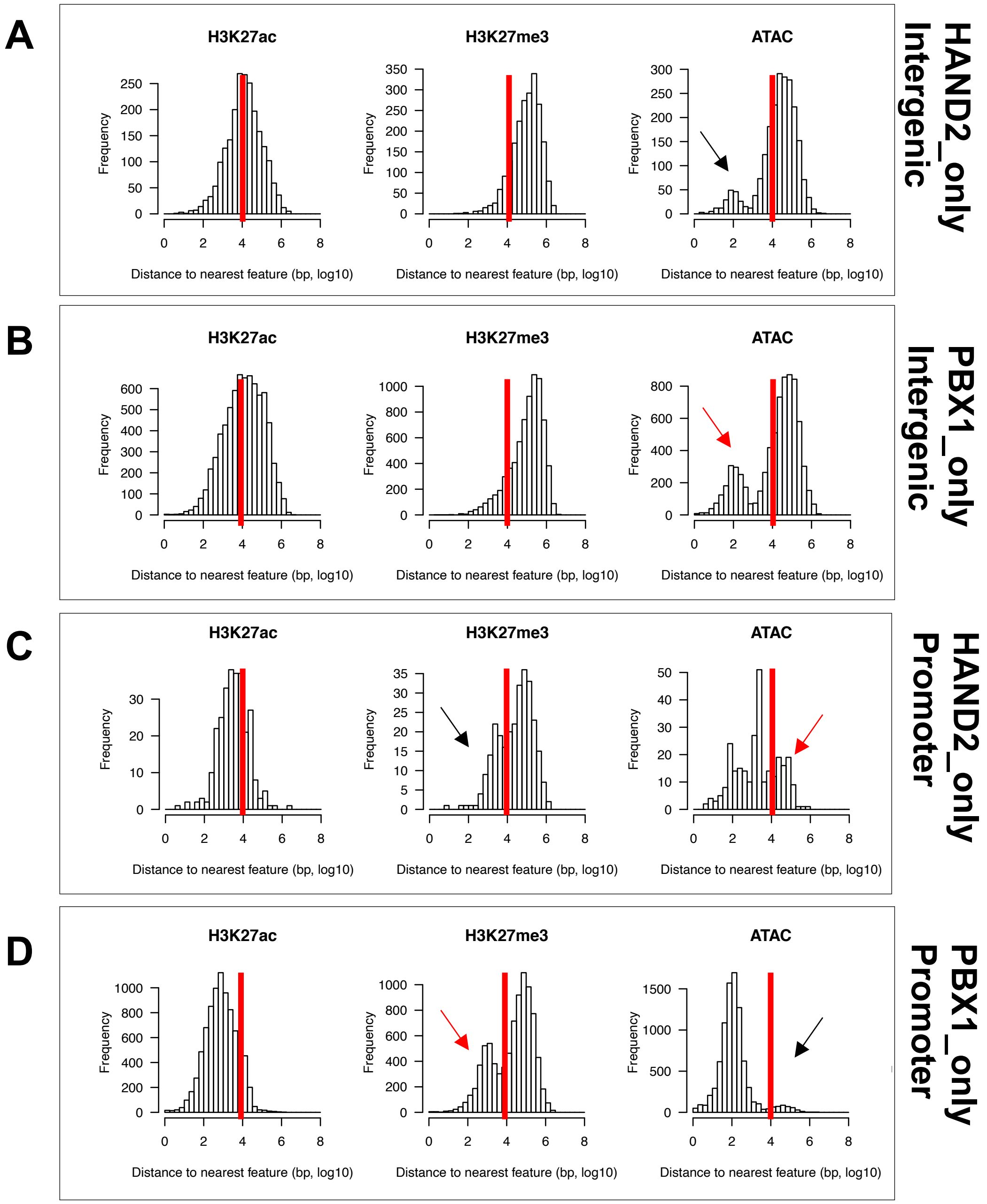

### Supplementary Figure 6

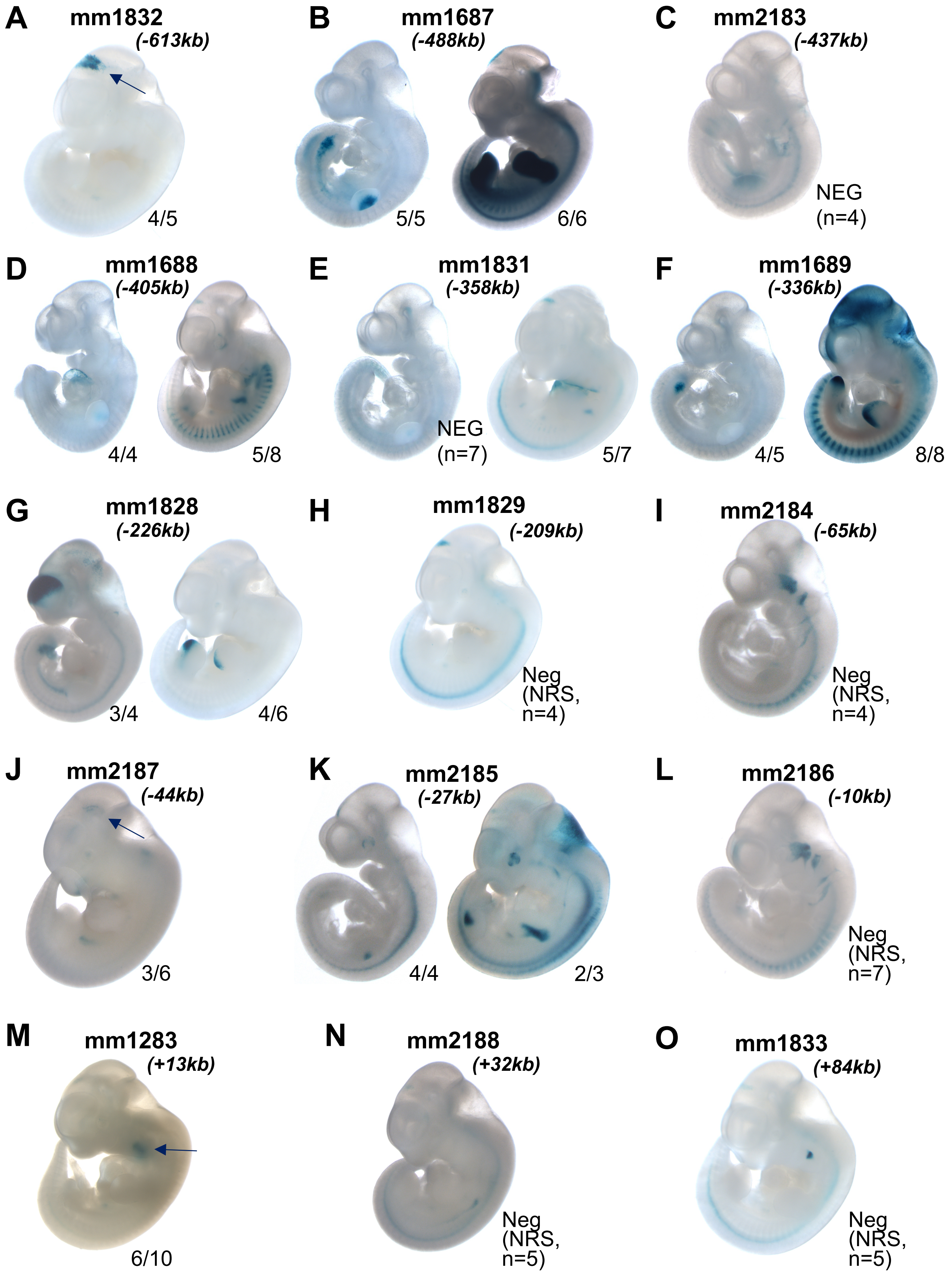

### Supplementary Figure 7

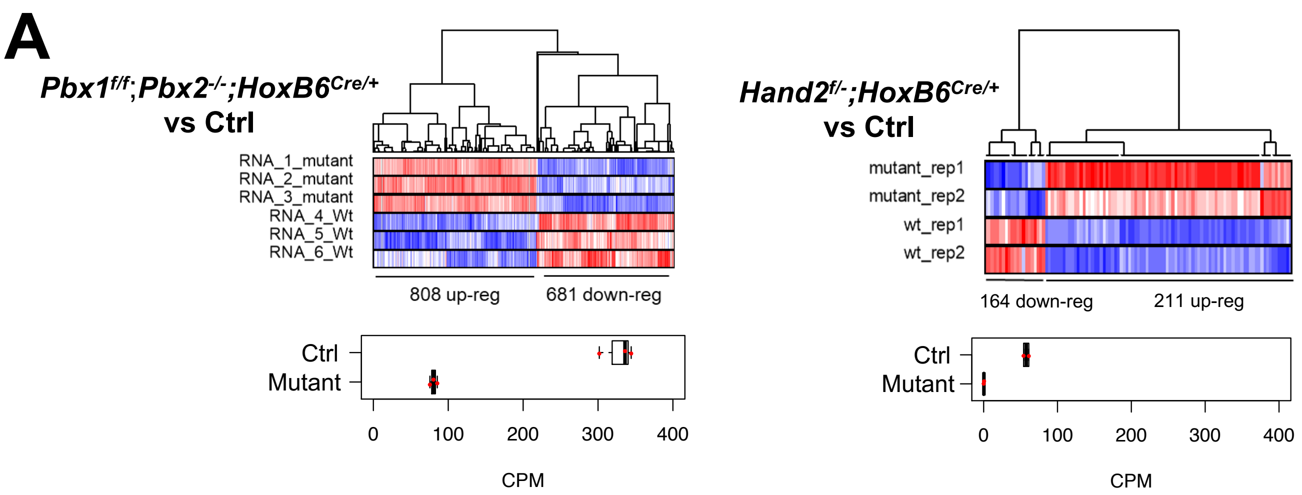


**B**

| | | | | |
|---|---|---|---|---|
| ***Pbx*** | **Up** | 808 | Not expressed in *Hand2* data | 25 |
| | | | *Pbx* only | 719 |
| | | | Both | 46 |
| | | | Discordant | 18 |
| | **Down** | 681 | Not expressed in *Hand2* data | 17 |
| | | | *Pbx* only | 614 |
| | | | Both | 37 |
| | | | Discordant | 13 |
| ***Hand2*** | **Up** | 211 | Not expressed in *Hand2* data | 15 |
| | | | *Hand2* only | 137 |
| | | | Both | 46 |
| | | | Discordant | 13 |
| | **Down** | 164 | Not expressed in *Hand2* data | 16 |
| | | | *Hand2* only | 93 |
| | | | Both | 37 |
| | | | Discordant | 18 |

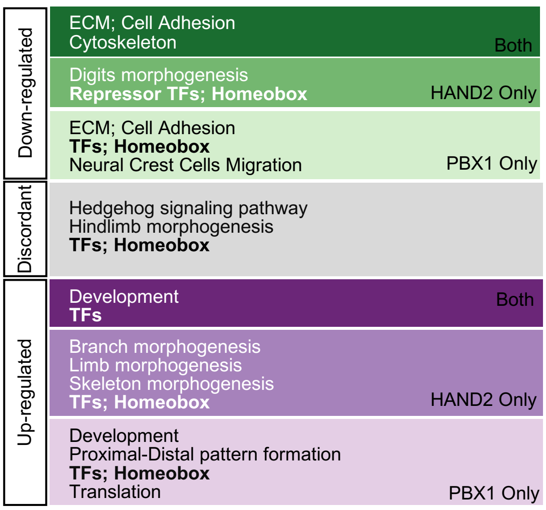


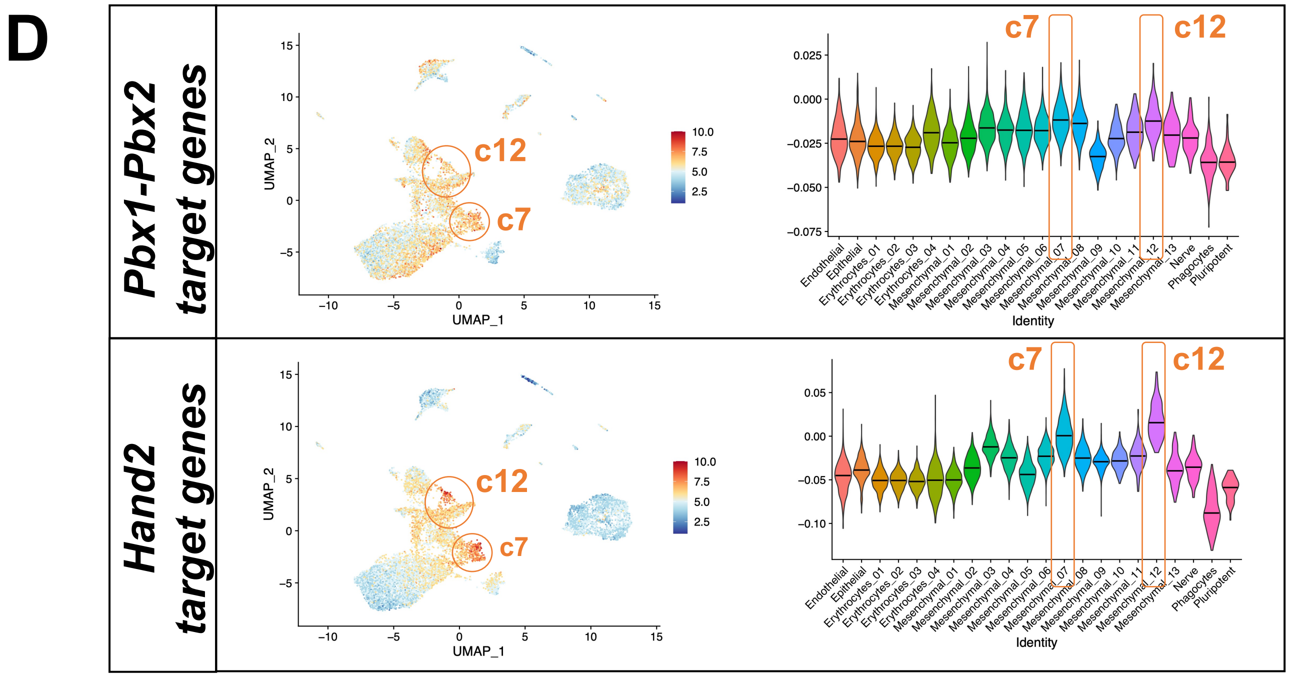

### Supplementary Figure 8

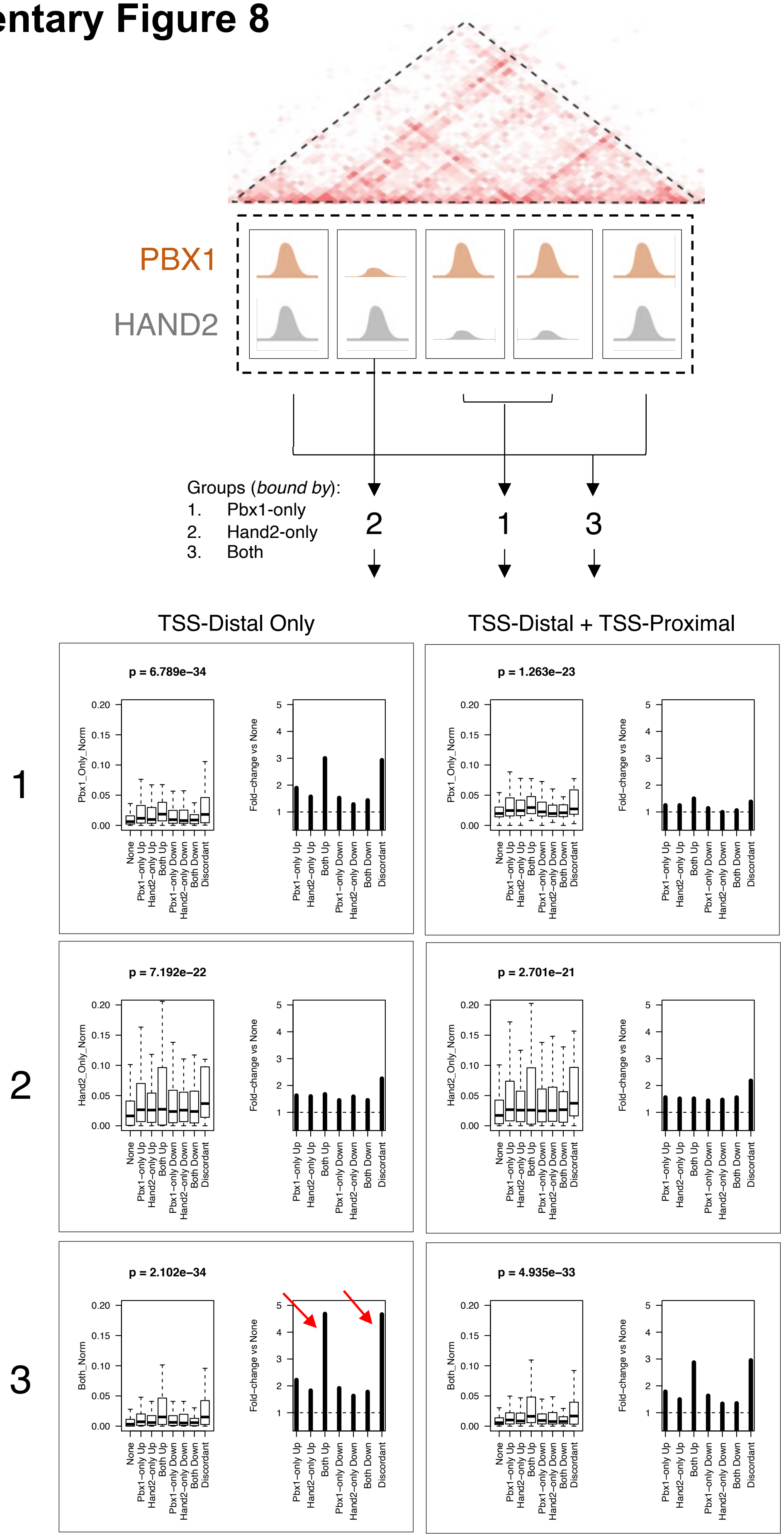

#### Supplementary Table 1

**A**

| Genotype | N | Gestational Days (E) | Skeletal Phenotype |
|---|---|---|---|
| Pbx1$^{f/+}$; Pbx2$^{+/-}$; Msx2Cre$^{+}$ | 15 | 10.5; 13.0; 14.5; P0 | no phenotype |
| Pbx1$^{f/+}$; Pbx2$^{-/-}$; Msx2Cre$^{+}$ | 8 | 10.5; 13.0; 14.5; P0 | no phenotype |
| Pbx1$^{f/f}$; Pbx2$^{+/-}$; Msx2Cre$^{+}$ | 12 | 10.5; 13.0; 14.5; P0 | no phenotype |
| Pbx1$^{f/f}$; Pbx2$^{-/-}$; Msx2Cre$^{+}$ | 8 | 10.5; 13.0; 14.5; P0 | no phenotype |
| Pbx1$^{f/-}$; Pbx2$^{+/-}$; Msx2Cre$^{+}$ | 6 | 13.5; P0 | no phenotype |
| Pbx1$^{f/-}$; Pbx2$^{-/-}$; Msx2Cre$^{+}$ | 5 | 13.5; P0 | no phenotype |

**B**

| Genotype | N | Gestational Days (E) | Skeletal Phenotype |
|---|---|---|---|
| *Pbx1$^{f/+}$; Pbx2$^{+/-}$; Hoxb6Cre$^{+}$* | 10 | 10.5; 13.5; 15.5; 17.5 | no phenotype |
| *Pbx1$^{f/+}$; Pbx2$^{-/-}$; Hoxb6Cre$^{+}$* | 11 | 10.5; 13.5; 15.5 | no phenotype |
| *Pbx1$^{f/f}$; Pbx2$^{+/-}$; Hoxb6Cre$^{+}$* | 10 | 10.5; 13.5; 15.5 | FL and HL defects |
| *Pbx1$^{f/f}$; Pbx2$^{-/-}$; Hoxb6Cre$^{+}$* | 8 | 10.5; 13.5; 15.5 | FL and HL defects |

#### Supplementary Table 2

Table attached as excel file

#### Supplementary Table 3

| Hand2 enhancers: elements tested | | | | |
|---|---|---|---|---|
| mm10 coordinates | Chr Start | Chr End | Distance to Hand2 TSS (in kb) | Vista ID |
| chr8 | 56706436 | 56707834 | -613 | mm1832 |
| chr8 | 56831311 | 56833826 | -488 | mm1687 |
| chr8 | 56883034 | 56884323 | -437 | mm2183 |
| chr8 | 56914551 | 56916754 | -405 | mm1688 |
| chr8 | 56961473 | 56963974 | -358 | mm1831 |
| chr8 | 56983040 | 56984959 | -336 | mm1689 |
| chr8 | 57093557 | 57095637 | -226 | mm1828 |
| chr8 | 57110606 | 57112872 | -209 | mm1829 |
| chr8 | 57254176 | 57256581 | -65 | mm2184 |
| chr8 | 57275519 | 57277110 | -44 | mm2187 |
| chr8 | 57292957 | 57294418 | -27 | mm2185 |
| chr8 | 57327753 | 57329581 | 7 | mm1284 |
| chr8 | 57330807 | 57332981 | 10 | mm2186 |
| chr8 | 57333128 | 57334891 | 13 | mm1283 |
| chr8 | 57353269 | 57354301 | 32 | mm2188 |
| chr8 | 57359264 | 57363084 | 40 | mm847 |
| chr8 | 57404702 | 57405866 | 84 | mm1833 |

#### Supplementary Table 4

Table attached as excel file

### Supplementary Table 5

| INTERSECTION PBX HL vs MF vs BA2 | Peaks | Total peaks per ChIPseq | Percentages |
|---|---|---|---|
| PBX HL | 6490 | 18423 | |
| PBX BA2 | 20045 | 31673 | |
| PBX MF | 7651 | 16850 | |
| Common PBX HL-BA2 | 4349 | | 54.35 |
| Common PBX HL-MF | 1920 | | 45.01 |
| Common PBX MF-BA2 | 1615 | | 43.20 |
| Common all | 5664 | | 33.61 |

| INTERSECTION PBX HL vs PBX BA2 vs HAND2 HL | Peaks | Total peaks per ChIPseq | Percentages |
|---|---|---|---|
| PBX HL | 7056 | 18425 | |
| PBX BA2 | 20179 | 31637 | |
| HAND2 HL | 2898 | 8405 | |
| Common PBX HL-BA2 | 7305 | | |
| Common PBX BA2-HAND2 HL | 1443 | | 13.13 |
| Common PBX HL-HAND2 HL | 1354 | | 48.35 |
| Common all | 2710 | | |

| INTERSECTION PBX HL vs PBX BA2 vs HOXA2 BA2 | Peaks | Total peaks per ChIPseq | Percentages |
|---|---|---|---|
| PBX HL | 8291 | 18426 | |
| PBX BA2 | 20921 | 31676 | |
| HOXA2 BA2 | 566 | 2306 | |
| Common PBX HL-BA2 | 9136 | | |
| Common PBX HL-HOXA2 BA2 | 121 | | 43.32 |
| Common PBX BA2-HOXA2 BA2 | 741 | | 70.21 |
| Common all | 878 | | |

| INTERSECTION HOXA2 BA2 vs HAND2 HL | Peaks | Total peaks per ChIPseq | Percentages |
|---|---|---|---|
| HOXA2 BA2 | 2030 | 2319 | |
| HAND2 HL | 8215 | 8504 | |
| Common | 289 | | 12.46 |

| INTERSECTION PBX HL vs PBX MF vs HAND2 HL | Peaks | Total peaks per ChIPseq | Percentages |
|---|---|---|---|
| PBX HL | 8049 | 18430 | |
| PBX MF | 8985 | 16851 | |
| HAND2 HL | 4093 | 8428 | |
| Common PBX HL-MF | 6328 | | |
| Common PBX HL-HAND2 HL | 2797 | | 48.09 |
| Common PBX MF-HAND2 HL | 282 | | 18.25 |
| Common all | 1256 | | |
